# Structural basis of antiphage defense by an ATPase-associated reverse transcriptase

**DOI:** 10.1101/2025.03.26.645336

**Authors:** Jerrin Thomas George, Nathaniel Burman, Royce A. Wilkinson, Senuri de Silva, Quynh McKelvey-Pham, Murat Buyukyoruk, Adelaide Dale, Hannah Landman, Ava Graham, Steven Z. DeLuca, Blake Wiedenheft

## Abstract

Reverse transcriptases (RTs) have well-established roles in the replication and spread of retroviruses and retrotransposons. However, recent evidence suggests that RTs have been conscripted by cells for diverse roles in antiviral defense. Here we determine structures of a type I-A retron, which explain how RNA, DNA, RT, HNH-nuclease and four molecules of an SMC-family ATPase assemble into a 364 kDa complex that provides phage defense. We show that phage-encoded nucleases trigger degradation of the retron-associated DNA, leading to disassembly of the retron and activation of the HNH nuclease. The HNH nuclease cleaves tRNA_Ser_, stalling protein synthesis and arresting viral replication. Taken together, these data reveal diverse and paradoxical roles for RTs in the perpetuation and elimination of genetic parasites.

## Introduction

Reverse Transcriptases (RTs) were originally discovered in Rous Sarcoma and murine leukemia viruses^1,2^. Over the next several decades, research on RTs suggested that these enzymes are primarily involved in the replication of selfish genetic elements, and occasionally domesticated for cellular functions (*e.g.,* telomerase and spliceosome)^3,4^. Initially, RTs were thought to be restricted to the eukaryotic domain of life^5-7^, but in 1989, three independent groups discovered RTs in bacteria that make multicopy single-stranded DNA (msDNA) using an RNA template ^8-10^. These systems were called retrons^6^, but the biological function of these systems remained enigmatic for almost 30 years^11^. Recently, these systems have been shown to provide bacteria with defense against phage infection^12,13^.

Retrons are diverse (*i.e.*, 13 types and several subtypes), but all retrons consist of an RT with two distinguishing sequence motifs (NAXXH and VTG), and a non-coding RNA (ncRNA) containing an inverted repeat and an invariant guanosine that acts as a primer for branched DNA synthesis^5,14,15^. Most retrons are associated with putative “effectors”, that are predicted to interact with cell membranes, cleave DNA, or degrade proteins^13,14,16,17^. To date, the only available retron structure is that of the type II-A3 (Eco1/Ec86) system^18-20^. These structures reveal an oligomeric assembly of retrons that sequester an N-glycosidase toxin, which is activated by phage-induced msDNA methylation^18,20^.

With the growing appreciation for the functional diversity of prokaryotic RTs in antiphage defense^13,21,22^, we were particularly intrigued by a class of RTs that are uniquely associated with Structural Maintenance of Chromosomes (SMC) ATPases^14^. SMC ATPases are conserved across all domains of life, perform critical functions in organizing higher-order chromosome structures, and facilitate DNA repair^23,24^. Intriguingly, SMC ATPases play critical roles in antiviral immunity in both eukaryotic (*e.g.*, human Smc5/6 and Rad50)^25-27^ and prokaryotic organisms (*e.g.*, PARIS^28-30^, Wadjet^31-33^, Lamassu^34-36^, and Gabija^36,37^). While RTs that synthesize extrachromosomal DNA and ATPases that recognize foreign DNA are independently recognized as components of antiviral defense, their unique association in type I-A retrons—sometimes called type II Septu^38^—raises fundamental questions about their functional synergy.

Here we clarify the functional interplay between RTs and SMC-family ATPases in type I-A retrons. We determine three different structures, which explain how RT-generated extrachromosomal msDNA serves as the structural core of a 364 kDa complex composed of the RT, a portion of the non-coding RNA, msDNA, two ATPase homodimers, and an HNH nuclease. Upon infection, a phage encoded nuclease cleaves the msDNA, triggering disassembly of the complex and activation of the HNH nuclease. The activated HNH selectively degrades cellular tRNA_Ser,_ arresting translation and stalling phage replication. Collectively, the structures, structure-guided mutants, phage challenge assays, and biochemical experiments presented here reveal how RT-generated extrachromosomal DNA—once viewed as a hallmark of selfish elements—acts as a crucial host factor that relies on SMC ATPases for antiviral defense.

## Results

### The type I-A retron system protects bacteria against phages

The type I-A retron system consists of a ncRNA, RT, msDNA, ATPase, and HNH nuclease (**Fig. 1a, Extended Data Fig. 1a,b**)^14^. Here we focus on a type I-A retron we identified in the FORC 82 strain of *Escherichia coli*. To determine if the *E. coli* FORC 82 retron provides phage defense, we cloned the operon along with ∼550 bases of the upstream DNA, which is expected to encode the ncRNA (**Fig. 1a**)^14^. The retron-containing plasmid or an empty vector control was transformed into *E. coli* MG1655, and plaque assays were performed using T3, T4, T5, and T7 phages. While the retron conferred defense against T3 and T5, no immunity was observed against T4 or T7 (**Fig. 1b, Extended Data Fig. 1c**).

**Figure 1.**
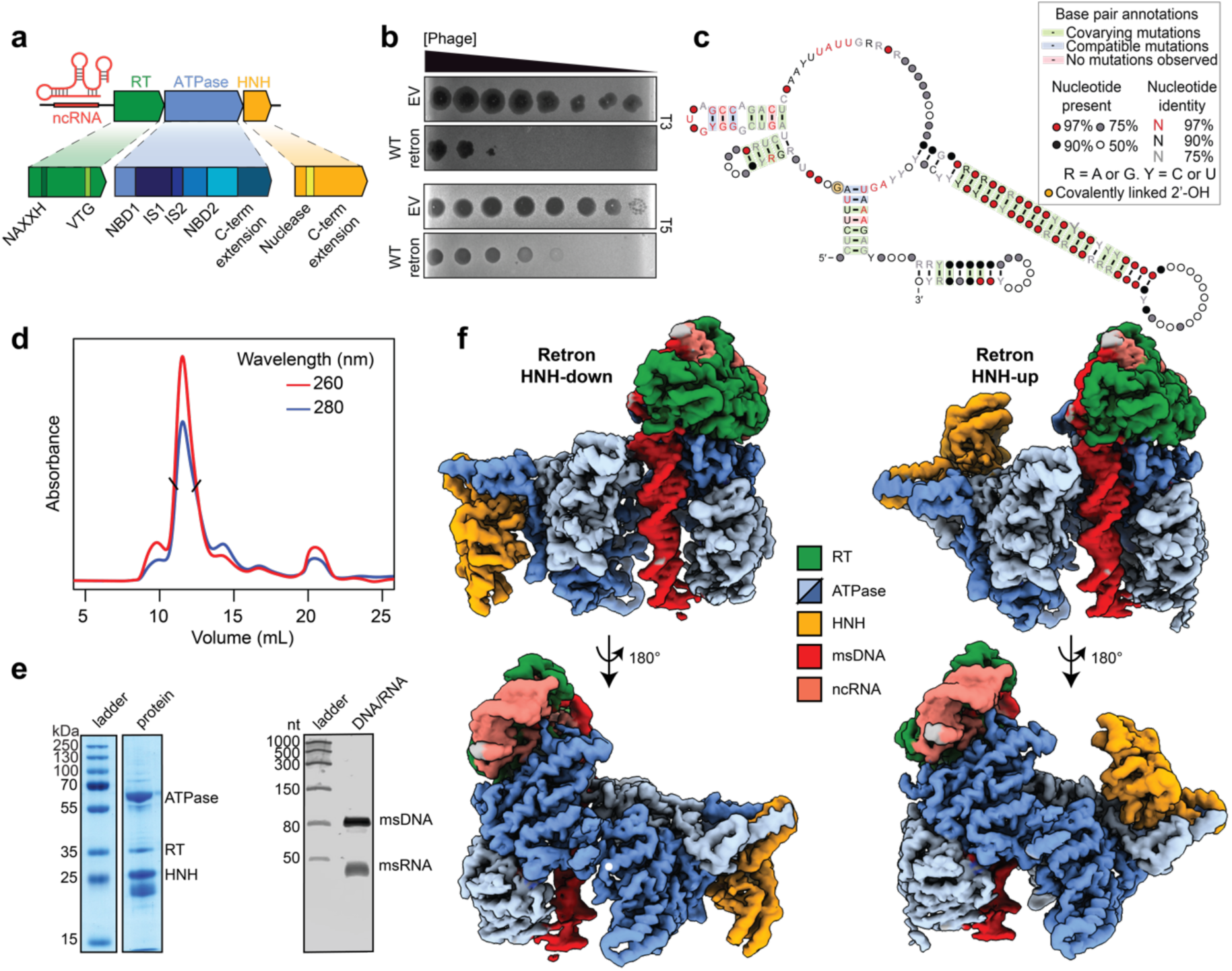
Phage defense and structures of a type I-A retron. **a,** Schematic of the type I-A retron operon. The retron encodes an RT, ncRNA, ATPase and HNH nuclease. Both the ATPase and HNH nuclease feature C-terminal extensions. **b,** Phage challenge assay demonstrate that the type I-A retron from *E. coli* protects against phages T3 and T5. The empty vector (EV) control exhibits no defense. Data shown is representative of *N* = 2 replicates. **c,** A covariance model for ncRNA from type I-A retrons generated from 70 sequences. **d,** Size exclusion chromatography (SEC) of the affinity-purified type I-A retron reveals a monodispersed peak with an estimated molecular weight of 350 kDa. The chromatogram is representative of *N* = 3 biological replicates. **e,** SDS-PAGE analysis of the main SEC peak reveals proteins corresponding to the ATPase, RT, and HNH. Nucleic acids resolved by denaturing PAGE show the ncRNA and msDNA. Data is representative of *N* = 3 biological replicates. **f,** Reconstructed density maps of the type I-A retron. HNH is bound asymmetrically in an up or down orientation to one of the ATPase homodimers.

To assess the role of each protein component in phage immunity, we introduced active site mutations in the ATPase (D387A), HNH nuclease (H71A), or the RT (YADD to YAAA). Mutations in the ATPase or HNH active sites eliminate retron-based phage defense, while attempts to clone mutation in the RT active site (YADD to YAAA) consistently failed, suggesting that these mutants are toxic (**Extended Data Fig. 1d**). Persistent attempts to clone the RT mutant resulted in the emergence of compensatory mutations in the ATPase (Q454K) or the HNH (G44S) and the ATPase (V73A) that collectively mitigate toxicity and rendered the defense system inactive (**Extended Data Fig. 1e, Supplementary Table 2**). These results demonstrate that all protein components of the *E. coli* FORC82 retron system are indispensable for defense, consistent with observations reported for retron Ec78^12,13^.

Given the involvement of multiple proteins, DNA, and RNA in the type I-A retron defense system, we hypothesized that the retron forms a multicomponent complex capable of recognizing phages and conferring immunity. To purify the complex, we first identified the 5’ and 3’ boundaries of the ncRNA by developing a covariation model built by aligning regions upstream of type I-A retron RTs (**Fig. 1c**). The model reveals a long stem-loop structure consistent with previous observations^14^. The predicted non-coding RNA and each of the protein coding genes were cloned into an expression vector with affinity tags on either the C-terminus of the HNH nuclease (HNH-Strep), the N-terminus of the RT (Strep-RT) or the N-terminus of the ATPase (Strep-ATPase). While the affinity-tagged proteins preserve retron-mediated immunity, only the tagged ATPase effectively pulls down all components of the retron (**Extended Data Fig. 1f,g**). The affinity-purified complex elutes from size-exclusion chromatography (SEC) as a monodispersed peak with an estimated molecular weight of 350 kDa. SDS-PAGE and mass spectrometry were used to identify all three protein components (*i.e.*, ATPases, RT, and HNH), while denaturing urea-PAGE and nucleases assays were used to identify the ∼90 nucleotide msDNA, and a ∼40 nucleotide ncRNA (**Fig. 1d,e, Extended Data Fig. 1h, Supplementary Table 3**). Our findings suggest that all components of the type I-A retron assemble into a multicomponent complex that is necessary for phage defense.

The purified complex was used for structure determination using cryo-EM (**Fig. 1f, Extended Data Fig. 2, Supplementary Table 1**). 2D classification revealed that a subset of particles contained an extra lobe of asymmetric density. This was reflected in initial 3D reconstructions which contained weak density for the additional domain. Focused 3D classification performed using a soft mask over the area of weak density produced three distinct structures with overall resolutions of 3.1 Å (**Fig. 1f, Extended Data Fig. 3**). Preliminary models of the complex were built by docking AlphaFold3^39^ predicted structures into the maps. This process revealed that the RT and a portion of the ncRNA comprise the “head” of the assembly, while the msDNA protrudes from the RT like a harpoon that extends all the way through the center of the complex. The HNH nuclease is asymmetrically bound to one of the two ATPase homodimers, in either an up or down configuration.

### Extrachromosomal DNA synthesized by RT is a substrate for the SMC ATPase

To understand how reverse transcription leads to formation of the retron, we compared an AlphaFold3-predicted structure of the ncRNA-RT complex to the experimentally determined structure of the retron complex (**Fig. 2a,b**). In contrast to the experimentally determined structure (*i.e*., post-assembly), the predicted structure represents an early intermediate. In both the predicted and experimentally determined structures, stem-loop 1 (SL-1) forms a molecular handle that interacts with the thumb domain of the RT and likely remains anchored to the RT^40,41^, while the RNA template is reeled through the active site during reverse transcription (**Fig. 2a-d, Extended Data Fig. 4a,c**).

**Figure 2.**
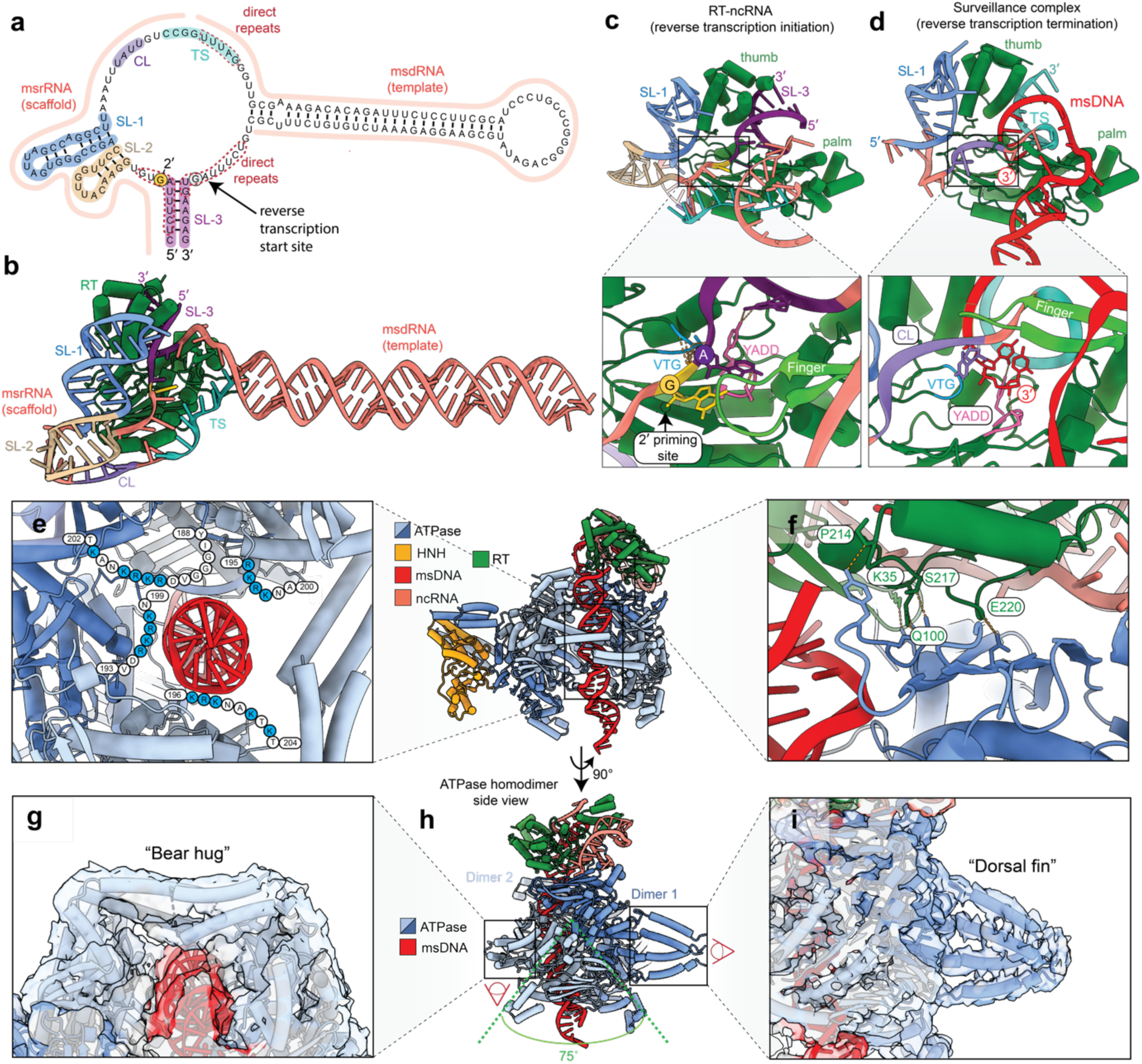
RT synthesizes a DNA ‘harpoon’ that stabilizes the complex. **a,** The ncRNA sequence of retron I-A from *E. coli* is mapped onto the covariance model shown in Fig. 1c. Conserved stem-loop motifs (SL-1, SL-2, and SL-3), the clamp (CL) and termination sequence (TS) are highlighted. **b,** AlphaFold3-predicted structure of the ncRNA-RT complex highlight features of the ncRNA identified in panel a. **c,** The ncRNA-RT structure prediction reveals that the RT thumb domain is trapped between SL-1 and SL-3, while SL-3 is positioned between the thumb and palm domains of the RT. The magnified region highlights the conserved VTG motif, which is predicted to make contacts with the DNA backbone, positioning the priming guanosine (G8, yellow) and adenosine 7 (A7, purple) near the active site (YADD, pink). **d,** Like the predicted structure of ncRNA-RT prior to msDNA synthesis (panel c), the experimentally determined structure of retron I-A reveals interactions between the RT thumb domain and SL-1. The duplex formed by the TS and the 3′ end of the cDNA is positioned between the thumb and palm domains, occupying the site previously occupied by SL-3 in panel c. The magnified region highlights the 3′ end of the cDNA contacting the YADD active site. **e,** The IS1 flexible loop of the SMC ATPase contains a series of basic residues that interact with the msDNA ‘harpoon’. **f,** The RT makes extensive interaction with one of ATPases (cyan). **g-i**, Coiled-coil domains in IS2 of the ATPase form distinct structural features on either side of the complex. **g,** On one side, the coiled-coil domains from opposing ATPases bear hug the DNA harpoon. **h,** On the opposite side, the IS2 coiled-coils form dorsal fin-like feature.

In the predicted ncRNA-RT structure, the 5′ and 3′ ends of the ncRNA form a conserved stem-loop (SL-3) that is positioned within a cleft of the RT formed by the thumb and palm domains (**Fig. 2a-c**)^42^. The prediction places the branching guanosine of the ncRNA, which serves as the primer for reverse transcription, near the VTG motif and the YADD active site (**Fig. 2c**)^14,15^. Reeling of the RNA template into the RT active site is anticipated to require unzipping the predicted stem of the ncRNA, and displacement of SL-3.

The clamp (CL) is a highly conserved feature of the ncRNA (**Fig. 2a**). This sequence is predicted to interact with the palm domain of the RT, prior to reverse transcription, while in the experimentally determined structure, the clamp is a linchpin that simultaneous contacts the 3′-end of the cDNA, the ATPase, and key domains of the RT, including the finger, palm, and VTG motif (**Extended Data Fig. 4d,e**). Collectively, these structural comparisons suggest that during reverse transcription, the 3′ end of the ncRNA is reeled through the active site and into the cleft formed by the thumb and palm domain (initially occupied by SL-3) while the SL-1 handle remains bound by the RT. Reeling of the template positions the 3′ end of the msDNA and the ncRNA termination sequence (TS) in close proximity, forming an RNA-DNA hybrid at the site previously occupied by SL-3 (**Extended Data Fig. 4d,e**).

The ATPase homodimers form an extensive network of electrostatic interactions with the msDNA (**Fig. 2e, Extended Data Fig. 5a**). Notably, a cluster of basic residues within insertion sequence 1 (IS1)—a conserved feature of SMC ATPases—is oriented towards the msDNA-binding cavity (**Fig. 2e**)^24^. However, IS1 is dynamic and density for IS1 is weak (**Movie 1**). While single-point mutants (K100A, K103A, K109A, K141A, N151A, N152A, K196A, or K201A) at residues involved in msDNA binding had no significant impact on phage defense (**Extended Data Fig. 5b**), multiple charge-swap mutations in IS1 were unclonable, suggesting toxicity (R195E, K196E, R197E, K198E, K201E, and K203E) (**Fig. 2e**). Similarly, structure-guided mutations that disrupt the interface between the RT and ATPase (RT residues: S217R, N219R and E220R) were also unclonable (**Fig. 2f**). Together, these findings underscore the critical role of msDNA and RT in orchestrating ATPase assembly, which is essential for the functional integrity of the retron.

SMC ATPases typically dimerize in the presence of ATP and play a central role in forming chromosomal structures that mediate DNA repair^23,24^. Insertion sequence 2 (IS2) is a conserved feature of SMC ATPases that form long coiled-coil domains, which wraps around genomic DNA^24,43^. In retrons, the ATPase dimers are offset by ∼75° relative to each other and coiled-coils from two adjacent ATPases encircle the extrachromosomal msDNA in an ATPase-mediated “bear hug” on one side of the complex and a “dorsal fin” shaped structure on the other (**Fig. 2g-i**). While the IS2-mediated bear hug obscures access to the msDNA, 3D variability analysis reveals a subset of particles with an alternate conformation, where the IS2 arms are disordered and the msDNA is exposed (**Movie 1**). Collectively, this reveals a conserved role for IS2 in binding chromosomal DNA during repair or msDNA during retron assembly.

### ATPase homodimers form a claw that grasps the HNH nuclease

The HNH nuclease is anchored to the retron complex via the C-terminal extension of the ATPase, which forms a triple helix (**Fig. 1a, 3a,b,d, Extended Data Fig. 1b**). The C-terminal triple helix from adjacent subunits form a claw-like structure that grips the C-terminal region of the HNH nuclease and structure guided mutations in ATPase (D522R, L523R, V526R, F529A, and K530D) result in a complete loss of defense (**Fig. 3a,b,d**). The nuclease is asymmetrically recruited to one side of the complex (*i.e.*, one pair of C-terminal extensions) in either an up or down orientation (**Fig. 1f**). Exclusion of the HNH from the opposing homodimer is partially explained by steric hindrance from the RT and SL-2 of the ncRNA, although clear density for SL-2 is not visible in our maps. In addition, three of the four ATPase nucleotide-binding domains (NBDs) include clear density for ATP, while the fourth NBD (NBD4) is reordered in a way that is incompatible with ATP binding (**Extended Data Fig. 6c,d**). This arrangement and exclusion of ATP from this NBD is consistent between the HNH-bound and HNH-free structures, demonstrating that the unique pose of NBD4 occurs during assembly rather than during HNH binding. Structural comparison with the type-I Septu ATPase (PDB: 8EEA) reveals a similar pattern of ATP exclusion, suggesting a conserved function in complex assembly, as opposed to HNH binding (**Extended Data Fig. 6e**)^44^.

**Figure 3.**
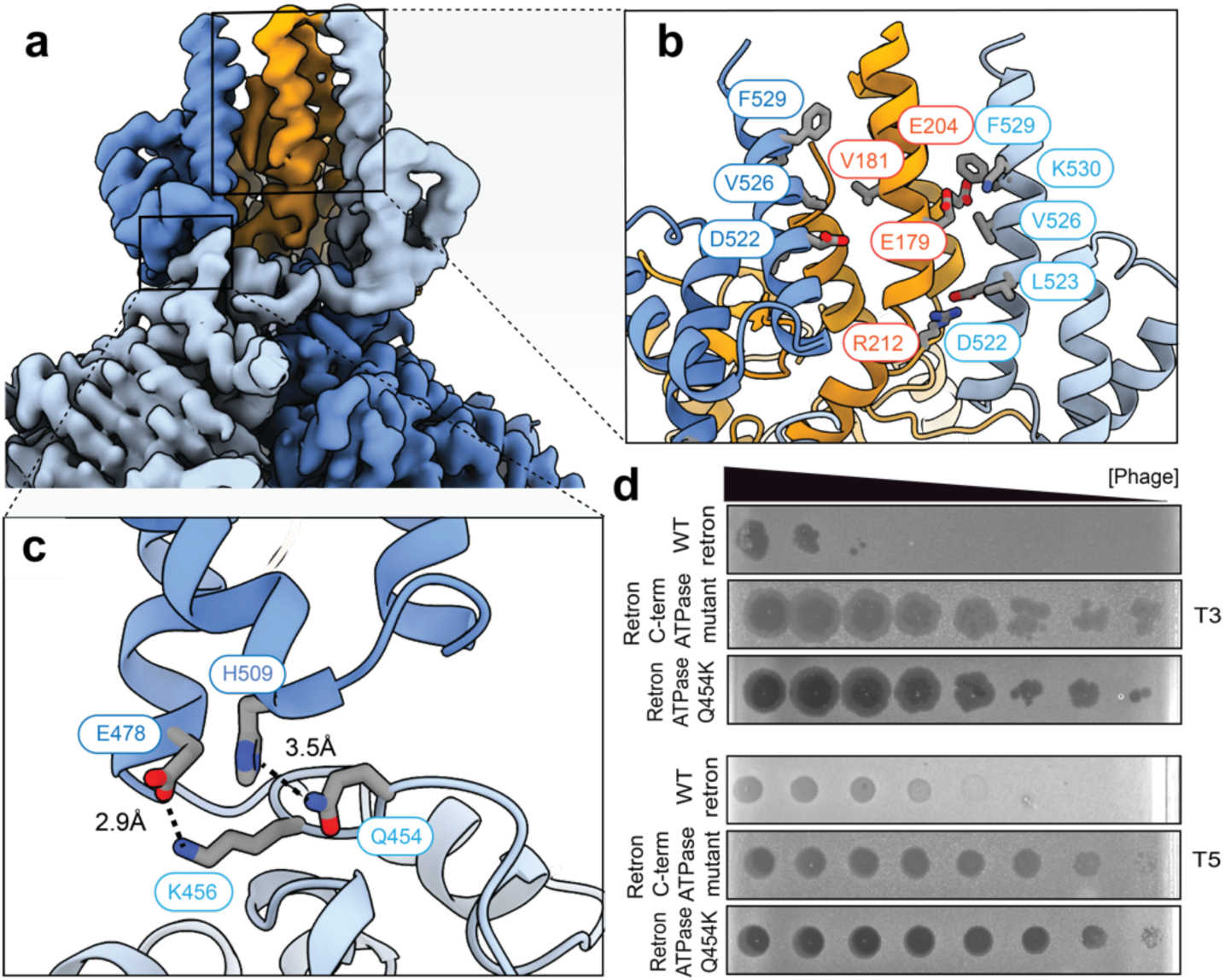
The C-terminal ATPase domain forms a claw-like structure that engages the HNH nuclease. **a,** Reconstructed density for the C-terminal ATPase domain interacting with the HNH nuclease. **b,** Key residues that mediate interactions between the ATPase and nuclease are highlighted. **c,** Q454 of the ATPase maintains structural integrity of the claw by stabilizing helices from the two oppositely oriented ATPase monomers. **d,** Phage challenge assay performed using a C-terminal mutant of the ATPase (D522R, F529A, K530E, L523R, V526R) or ATPase mutant Q454K. Data shown is representative of *N = 2* replicates.

While mutations in the RT active site render the retron toxic, a spontaneous mutation in the ATPase (Q454K) renders the RT active site mutants non-toxic. Q454 is conserved across type I-A retron ATPase, and the structure explains a role for this residue in stabilizing the claw-like structure (**Fig. 3c**). Although Q454 does not directly interact with the HNH nuclease, the Q to K mutation leads to a complete loss of defense, highlighting the importance of the claw-like structure in recruitment of the HNH (**Fig. 3d**).

### The msDNA is a phage sensor

Building on our structural understanding, we hypothesized that retron activation is triggered by specific components of the infecting phage. To test this hypothesis, we screened for phage mutants that escape retron-mediated defense. Cells expressing the retron defense system were infected with phages T3 and T5. Three T3 escape mutants were isolated, but no escapers were recovered for T5 despite repeated attempts (**Extended Data Fig. 7a**). Whole-genome sequencing of the T3 escape mutants revealed mutations in four phage genes (serine/threonine kinase, phage tail fiber, phage tail protein, and an exonuclease), as well as two additional mutations in non-coding regions (**Fig. 4a**). While serine/threonine kinase was toxic by itself, the other genes were expressed either alone or in retron-containing cells. Co-expression of the exonuclease with the retron severely impaired bacterial growth, whereas no growth defect was observed when the exonuclease was expressed alone or with a retron containing an inactive HNH nuclease (H71A) (**Fig. 4b,c**). These results indicate that T3 exonuclease triggers retron-mediated immunity.

**Figure 4.**
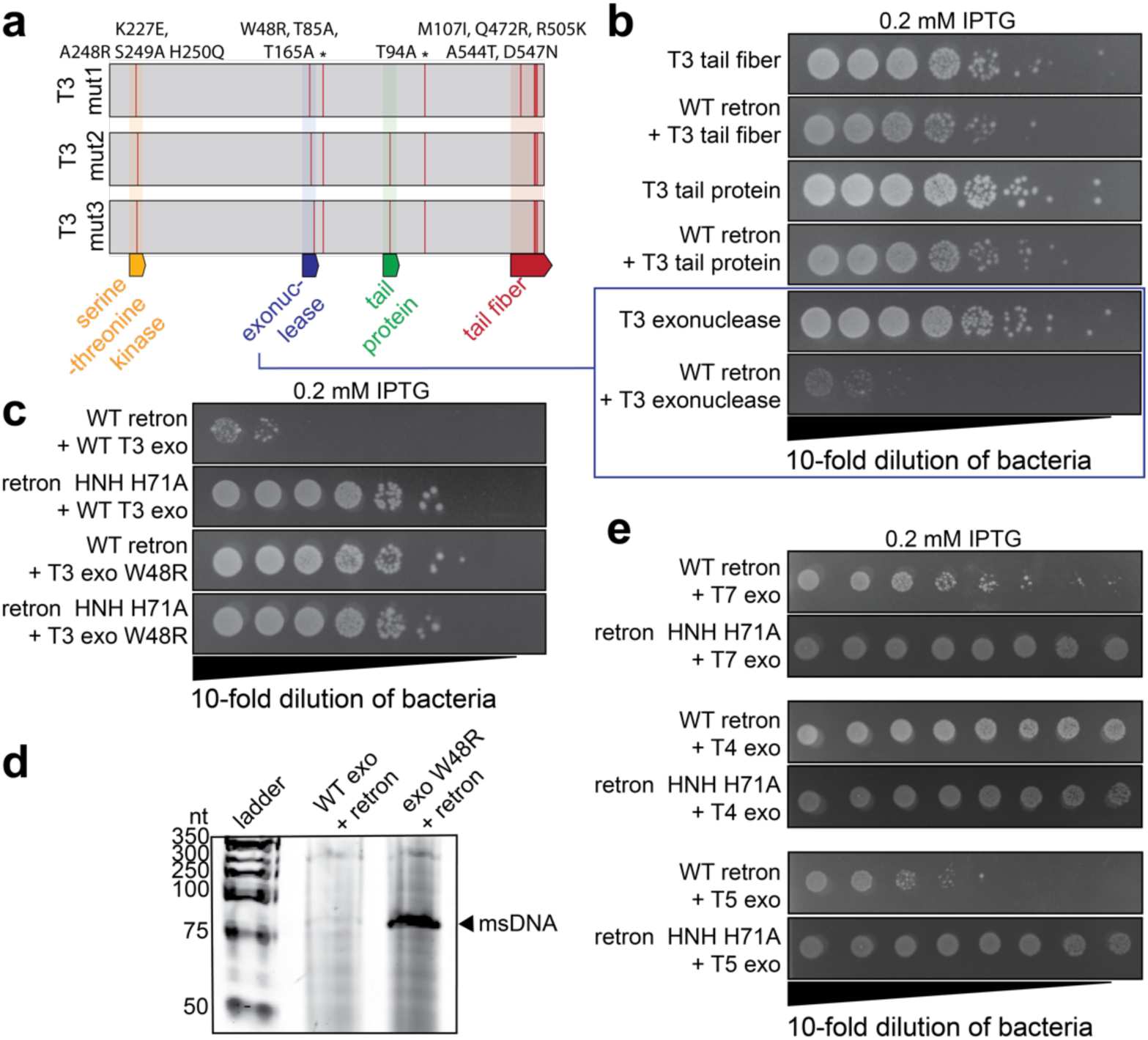
T3 exonuclease activates the retron defense system by cleaving msDNA. **a,** Genome sequencing of three T3 escape mutants reveals the locations of mutations (red lines). Mutations in non-coding regions are marked with asterisks. **b,** Dilution series of *E. coli* cells shows the effects of expressing phage genes alone or in combination with the retron. **c,** Dilution series of *E. coli* cells expressing retron mutants reveals that mutations in the HNH nuclease (control) or the phage exonuclease, both abolish cell toxicity. **d,** Urea-PAGE analysis confirms the presence or absence of msDNA in samples co-expressing WT or mutant exonuclease with the retron. **e,** Dilution series of *E. coli* cells expressing exonuclease from phages T7, T4, or T5, with WT or mutant (H71A) retron. Data shown in panel b, c, and e are representative of *N = 2* biological replicates.

An AlphaFold3 predicted structure of the T3 exonuclease reveals that one of the escaper mutations (W48R) is on a β-hairpin ∼34 Å from the exonuclease active site (**Extended Data Fig. 7d**). Co-expression of the T3 W48R exonuclease mutant with the retron did not result in cell growth arrest, suggesting that retron activation is dependent on specific structural features beyond the active site (**Fig. 4c**). Mapping the remaining escaper mutations on the exonuclease structure revealed that none of the mutations occurred at the active site (**Extended Data Fig. 7d**). This suggests that the exonuclease activity is not required for retron activation, or this activity is essential for phage replication and cannot tolerate mutations that disrupt its function. To determine whether the exonuclease directly degrades msDNA, acts indirectly through other cellular nucleases, or has no impact on msDNA integrity, we mutated the exonuclease active site (D162K) and co-expressed the mutant or the wild-type (WT) exonuclease in retron containing cells. Co-expression of the T3 exonuclease active site mutant (D162K) or T3 phage escaper mutant (W48R) with retron results in no growth arrest and no degradation of the msDNA, while the WT exonuclease results in both growth arrest and msDNA degradation (**Fig. 4c,d, Extended Data Fig. 7b,c**). Collectively, this data demonstrates that the T3 exonuclease triggers retron activation, that residues outside of the active site are necessary for retron-mediated recognition of the trigger, and that the nuclease active site is necessary for degrading the msDNA.

To better understand the mechanism of retron activation by the T3 exonuclease, we identified and compared related nucleases using a combination of AlphaFold3^39^ and FoldSeek^45^ (**Extended Data Fig. 7d,e**). T3 exonuclease is structurally similar to the T4, T5, and T7 flap nucleases that have been proposed to remove Okazaki fragments (**Extended Data Fig. 7e**)^46-49^. To determine if the other phage encoded nucleases activate the retron, we co-expressed the T4, T5, and T7 nucleases with the retron (**Fig. 4f**). Nucleases from T3 and T5 (i.e., D15) trigger retron-mediated growth arrest, while the T4 and T7 nucleases have little impact on cell growth (**Fig. 4e, Extended Data Fig. 7e**). These experiments, performed by expressing only the viral nucleases, are consistent with results from plaques assays performed with the T3, T4, T5 and T7 phages (**Fig. 1b, Extended Data Fig. 1c**), verifying that retron-mediated protection from T3 and T5 phages is due to nuclease-mediated activation of the retron.

### Retron activation triggers translation arrest

The type I Septu defense system, composed of an ATPase and an HNH nuclease, degrades viral DNA during infection^44^. Growth arrest following type I-A retron activation, along with structural similarity between the retron and Septu HNH nucleases, led us to hypothesize that the retron would rely on a similar mechanism (**Extended Data Fig. 6e**). To test this hypothesis, we repeated experiments done for Septu by inducing *E. coli* cells expressing either the WT or an HNH-dead mutant (H71A) retron along with the T3 exonuclease and analyzed the cellular DNA using fluorescence microscopy (**Fig. 5a**). Rather than DNA degradation, we observed nucleoid compaction and propidium iodide exclusion (**Fig. 5a, Extended Data Fig. 8**). In bacteria, transcription, translation, and membrane insertion of proteins are tightly coupled through a process known as transertion, and disruption of translation has been shown to induce DNA compaction^50-52^. Collectively, these data suggest that retron-mediated growth arrest is due to translational inhibition.

**Figure 5.**
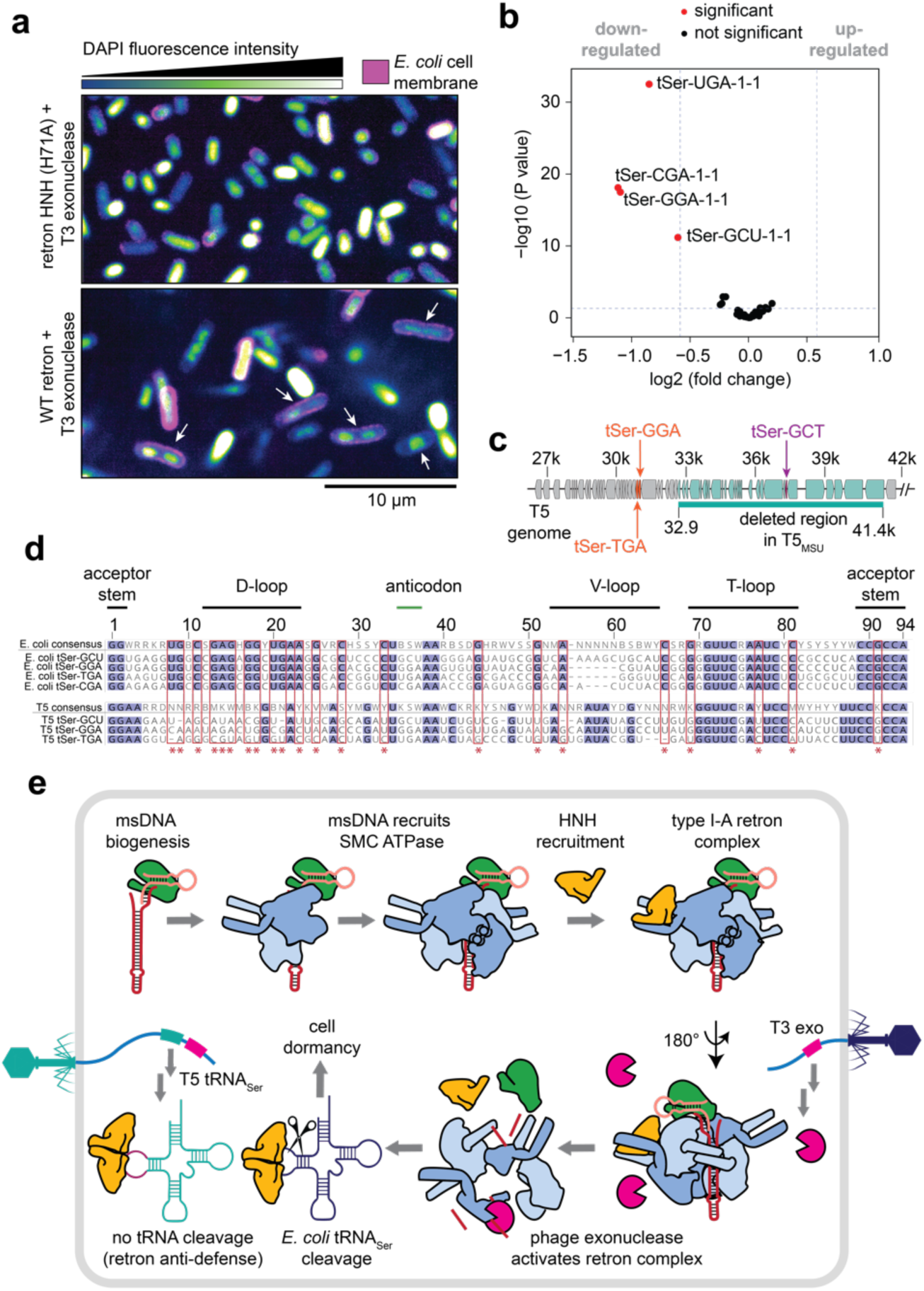
HNH nuclease degrades *E. coli* tRNA_Ser_. **a,** Fluorescence microscopy images for WT or HNH-inactive (H71A) retron in the presence of T3 exonuclease. Cell membrane is labeled with WGR Oregon (violet). DNA is labeled with DAPI (blue). Arrows indicate cells with DNA compacted away from the cell membrane. Images representative of *N* = 2 independent biological replicates. **b,** Volcano plot comparing tRNA abundance between cells expressing WT retron and T3 exonuclease versus those expressing an HNH-inactive retron and exonuclease. The *p* value is calculated using a negative binomial test. Data shown is representative of *N* = 3 independent biological replicates. **c,** WT T5 sequence with the deleted region in T5_MSU_ highlighted in cyan. The location of the phage tRNA_Ser_ are indicated. **d,** Alignment of *E. coli* and T5 tRNA_Ser_. Red boxes and asterisk indicate residues conserved in *E. coli* tRNA_Ser_ but different in T5 tRNA_Ser_ **e,** Proposed model for antiphage defense by the ATPase-associated RT. The RT reverse-transcribes a ncRNA into an extrachromosomal DNA that recruits SMC ATPases. The HNH toxin is recruited. The phage flap endonuclease degrades the msDNA, releasing the HNH, which depletes *E. coli* tRNA_Ser_.

During the course of this work, Azam *et al.* reported that expression of a related ATPase and HNH nuclease isolated from the type I-A retron (Ec78) from strain 102598 of *E. coli*, led to the degradation of bacterial tRNA_Tyr_ (**Extended Data Fig. 1**)^53^. Additionally, they showed that T5 phage encodes a tRNA_Tyr_ resistant to cleavage by the retron system. Inspired by these findings and results from our microscopy, we performed tRNA sequencing of cells expressing the T3 exonuclease with either the WT retron or HNH-inactive (H71A) control. This analysis reveals a significant depletion of all four serine tRNAs encoded in *E. coli* MG1655 (**Fig. 5b, Extended Data Fig. 9a,b**). However, the T5 genome also includes tRNAs for serine, which would be anticipated to complement tRNAs depleted by the activated retron. To verify the T5 genome, we sequenced and compared the T5 strain we were using to the WT T5 genome in NCBI (T5_WT_). The T5 strain used in these experiments (designated T5_MSU_) includes a 8.6 kb deletion within the tRNA-rich genomic region (**Fig. 5c**)^54^. This deletion eliminates 10 of the 25 tRNAs including a tRNA_Ser_ containing a GCT anticodon (tRNA_Ser_-GCT), which decodes a highly represented codon in both the T5 and *E. coli* genomes (**Extended Data Fig. 9c**). The deletion of T5 encoded tRNAs may occur spontaneously during serial passage of the virus in the absence of selective pressures (e.g., immune system), and the deletion of tRNA_Ser_-GCT renders the mutant strain (T5_MSU_) sensitive to the immune system.

To investigate how viral tRNAs avoid retron-mediated elimination, we aligned *E. coli* tRNA_Tyr_ and tRNA_Ser_ sequences to analogous tRNAs encoded in the T5_WT_ genome (**Fig. 5d, Extended Data Fig. 10**). Notably, most differences between the phage and *E. coli* tRNAs cluster in or around the D-loop, a region recognized by aminoacyl-tRNA synthetases^55^. These findings suggest that these two HNH nucleases target the D-loop in specific tRNAs, and that T5 phages evade retron-mediated immunity by encoding tRNAs with D-loop mutations. To determine if D-loop variation in tRNAs is a more general theme, we aligned *E. coli* tRNAs to each of the corresponding tRNAs in T5. All 23 tRNAs in the T5 genome contain D-loop mutations relative to analogous isoaccepters in *E. coli*. However, mutations were not restricted to the D-loop, which is consistent with a wider spectrum of nucleases that target tRNA degradation and modification^28,30,56,57^.

## Discussion

The competing selfish interest of viruses and their hosts results in a biological tit-for-tat that necessitates perpetual innovation to avoid extinction^58-60^. As part of this process, genes with one function are often conscripted, repurposed, or swapped between the competing interests in ways that (temporarily) enhance fitness^61,62^. Here we determine structures, perform structure-guided mutations, phage challenge assays, and biochemical experiments that collectively explain how bacteria and archaea use RTs to generate extrachromosomal DNA, that serves as a scaffold to recruit SMC ATPases and HNH nucleases for antiviral defense. The experimentally determined structure and structure predictions add new insight to the process of reverse transcription, and explain how the RT links the ncRNA and msDNA to a pair of SMC ATPase homodimers (**Fig. 5e**). Two insertion sequences (IS1 and IS2), typical of SMC ATPases, make extensive interactions with the msDNA, while C-terminal extensions on two adjacent ATPases form a claw-like structure that engages the HNH nuclease, while analogous residues on the other pair of ATPases are disordered (**Fig. 1**). The rational and mechanism for only recruiting one HNH per retron remains unclear.

The C-terminal claw of the ATPase and the HNH are structurally similar to that previously reported for Septu (RMSD = 1.1 Å across 56 atom pairs, and 2.7 Å across all 84 pairs) (**Extended Data Fig. 6a,b**)^44^. However, Septu systems do not involve RTs, the HNH is reported to be a DNase rather than an RNase, and the viral activator is unknown^44^. The HNH structure reported here is nearly identical to the HNH from Septu (RMSD = 1.1 Å across 101 pruned atom pairs), so it remains unclear how subtle structural differences result in distinct biological outcomes. Nevertheless, the ability of retron-associated HNHs to cleave tRNAs makes them unique among HNH family nucleases that are frequently associated with DNA cleavage^63^.

Phage-encoded nucleases trigger msDNA degradation, but the mechanism of degradation remains unclear. According to the structure, the msDNA and/or ncRNA are vulnerable to nuclease activity at the head (SL1, SL2 and/or SL3), the body (shaft of the msDNA), or poorly ordered tip of the msDNA, which includes 12 unmodeled bases that extend beyond the ATPases. Initially we suspected the tip, because it is most exposed and because the covariance model predicts high variability in this region, which could reflect a mechanism for identifying different phage triggers. However, truncation of the loop had no impact on defense (**Extended Data Fig. 7f**), suggesting that the tip of the harpoon may not be directly involved in trigger detection. Multiple efforts to use AlphaFold3 to predict a binding site for the trigger never resulted in a high confidence prediction, but most of the models positioned the T3 exonuclease near the head of the complex. Verification of these models will require further structural studies.

Recently, Azam *et al.* reported that co-expressing the ATPase and HNH nuclease from a type I-A retron in *E. coli* (Ec78) depletes cellular tRNA_Tyr_^53^, whereas we reveal that the retron from *E. coli* FORC 82 depletes all serine tRNAs. While this divergence in tRNA specificity suggests that different retron systems have evolved distinct substrate preferences, our findings suggest that both systems converge on targeting the D-loop, indicating a conserved recognition strategy. This model parallels what we and others observed for the TOPRIM nuclease in the PARIS defense system, where distinct tRNAs are cleaved at the same structural location and virally encoded tRNAs antagonize these systems^28,57^. Overall, these findings highlight the paradoxical role of RTs in viral propagation and antiviral defense, the functional plasticity of SMC ATPases (e.g., DNA repair and immunity), and illuminate a growing appreciation for the role of tRNAs as biological “double agents” that are sculpted by the selective pressures of genetic conflict.

## Methods

### Plasmid construction

Phage challenge, toxicity assays, tRNA sequencing and msDNA cleavage analysis were performed in *E. coli* MG1655, with the WT retron operon from *E. coli* FORC 82 cloned into a pACYCDuet-1 backbone with native promoters. Overexpression and large-scale protein pull-down experiments were performed using T7 promoters in a pRSF backbone, with an N-terminal Strep-tag on the ATPase. The phage triggers were cloned into pCDFDuet-1. Protein expression and pull-down assays were conducted in *E. coli* BL21 AI cells. The retron operon from *E. coli* FORC 82 was synthesized as a gene fragment by GenScript. The triggers were amplified and cloned from respective phages.

All primers used for cloning were obtained from Eurofins Genomics. Cloning was performed in either DH5α or NEB Turbo competent *E. coli* cells, and plasmids were extracted using the Zymo ZR Plasmid Miniprep Kit. The sequence of each cloned construct was verified by whole plasmid sequencing using Oxford Nanopore (Plasmidsaurus).

*E. coli* cultures were grown on LB agar plates or in LB broth. Constructs with the pACYCDuet-1 backbone were maintained with 25 μg/mL chloramphenicol, pRSF constructs with 50 μg/mL kanamycin, and pCDFDuet-1 constructs with 100 μg/mL Streptomycin unless otherwise specified. All plasmids used in this study are provided in Supplementary Table 2.

### Phage challenge assay

Plaque assays were performed using high phage titers (10⁸ PFU/mL) of T3, T4, T5, and T7. *E. coli* MG1655 cells transformed with plasmids for the WT retron, empty vector (pACYCDuet-1), or mutant retrons were grown to an OD600 of 0.3–0.4 in LB media supplemented with chloramphenicol (34 µg/mL). The cells were added to 0.6% agarose in LB (top layer) and the suspension was poured onto plates containing 3.2% LB agar (bottom agar). Both the top layer and the bottom agar were supplemented with chloramphenicol and 2 mM CaCl₂. Once the top layer solidified, 2.5 µL of 10-fold serial dilutions of phage stocks were spotted on the lawn of *E. coli* cells. The plates were then incubated overnight at 37 °C.

### Covariance model and sequence comparison for ncRNA and tRNA

Retron type I-A containing genomes^14^ were downloaded from NCBI (*N* = 70). 1 kb upstream of each RTs was extracted using a custom Python script and aligned using MAFFTv7.520^64^ with the following parameters “*--maxiterate 1000 --localpair.*” Alignments were manually curated to determine the exact boundaries of the ncRNA (∼250 bp region). The conserved region was extracted and realigned with mLocARNA v2.0.0^65^ using “*--stockholm --concensus-structure alifold*.” The resulting alignment was used to generate a covariance model that was visualized with R2R v1.0.7^66^.

*E. coli* MG1655 and WT T5 genomes were downloaded from NCBI database. tRNAscan-SE web server^67^ was used to predict T5 tRNAs. mLocARNA v2.0.0^65^ was used to align tRNA sequences to generate sequence alignments and covariance model.

### Recombinant expression and purification of the retron complex

The N-terminally Strep-tagged ATPase along with the other type I-A retron components, were co-expressed using an inducible T7 promoter in *E. coli* BL21 AI cells. Cultures were grown in LB media containing 50 μg/mL kanamycin at 37 °C until OD600 ∼0.5, then induced with 0.2 mM Isopropyl β-D-1-thiogalactopyranoside (IPTG) and 0.1% arabinose and grown for 16 h at 22°C. Cells were harvested by centrifugation at 3,000 × *g* for 10 min at 4°C, then lysed by sonication in lysis buffer (20 mM Tris pH 7.5, 200 mM KCl, 2 mM MgCl₂, 1 mM DTT). The lysate was clarified by centrifuging at 10,000 × *g* for 25 min, and the supernatant was passed through a 5 mL StrepTactin column (IBA Lifescence). After washing with the 20-column volumes of the lysis buffer, bound proteins were eluted using lysis buffer supplemented with 2.5 mM desthiobiotin. The retron was further purified using a Superdex 200 10/300 column equilibrated with 20 mM Tris pH 7.5, 200 mM KCl, 2 mM MgCl₂, 1 mM DTT, 5% glycerol. Fractions containing the purified retron components were analyzed by 12% SDS-PAGE to resolve protein components, while 15% urea-PAGE was used to visualize ncRNA and msDNA. The fraction containing the retron complex was further concentrated and used for vitrification

### Cryo-EM sample preparation

C-flat copper grids with a mesh size of 300 and R1.2/1.3 hole spacing were glow discharged using a Pelco EsiGlow at 15 mA for 45 seconds. 3 µL of 4 µM retron was applied to prepared grids in a Vitrobot Mk IV (ThermoFischer) set to 100% humidity and 4°C. Grids were subjected to double-sided blotting with a force of 5 for 5s before plunge-freezing in liquid ethane. Clipped grids were loaded into autogrid boxes and stored in liquid nitrogen before imaging.

### Cryo-EM data collection

Data were collected on a Talos Arctica (Thermo Fischer) equipped with a Gatan K3 direct electron detector at Montana State University’s Cryo-EM core facility using SerialEM^68^ under the control of SmartScope^69^ for automated data collection. 15,594 micrographs were collected with a pixel size of 0.9061Å and apparent magnification of 45,000X. The total dose per exposure was calculated as 56.69 electrons with 37 subframes per exposure (1.53el/px/frame).

### Cryo-EM data processing

After filtering micrographs by CTF-fit <8 Å resolution and total motion < 100 px, 13,695 exposures were subjected to blob picking in cryoSPARC^70^ Live (60-200Å diameter). 1,576,065 blobs were extracted using a box size of 512px (binx2) and sorted by 2D classification to yield a stack of 358,381 selected particles. From these particles, a *de novo* template volume was generated which contained 245,457 particles and refined to 3.7 Å. Using this volume as a template for particle picking, 7,373,097 particles were extracted with a box size of 512 px (binx4) and subjected to 2D classification with a 200 Å mask. 1,321,836 particles were selected and fed into a 3-class *ab initio* reconstruction. One class was identified containing 1,244,961 particles that appeared to correspond to the retron complex but exhibited compositional heterogeneity in regions of density corresponding to the HNH. Using a soft-padded mask directed towards the HNH, focused 3D classification yielded three distinct particle classes that varied based on the orientation (up or down), or absence of the HNH subunit. Un-binned particles were then re-extracted with a box size of 480px for non-uniform refinement. After reference-based motion correction, we carried out a final round of 2D classification to remove remaining junk. Final reconstructions were obtained from non-uniform refinements with several options enabled including minimizing over per particle scale, correcting for per-particle defocus and estimating the exposure group parameters Cs, Anisotropic Magnification, and tetrafoil. Phenix’s^71^ half-map based sharpening was used to aid in model building..

### Model building

An initial model was established by docking AlphaFold3 predicted structures into the density^39^. After rigid body fitting the individual chains using the “fit in map” command in ChimeraX^70^, each chain was equilibrated in ISOLDE^72^ to improve the initial model. For each density map, the corresponding maps and models were provided to Phenix^71^ for real-space refinement. Per residue CC scores were used to guide the trimming of flexible loops and regions of poor density. Sidechains not visible in the map were removed. Maps were sharpened using Phenix’s implementation of anisotropic half-map sharpening to aid in modeling difficult regions. Additionally, 3DVA volumes were used to inform the position of the ATPase coiled-coil domains^73^. While these features were not included in the mask during the final iterations of high-resolution refinement, they are clearly visible in lower-resolution refinements of the same particle stacks used to generate the maps submitted to EMDB (Supplementary Table 1). As such, we opted to include the coiled coil domain in the PDBs, which helps to explain how access to the msDNA is gated by ATPase subunits, as well as highlight the role of the coiled coil in the tetrameric assembly of ATPases.

### Phage escaper screening and genome sequencing

Phage escaper assay were performed as described in Avigail et al^17^. Briefly, *E. coli* MG1655 cells with or without the retron were grown in LB media at 37°C to mid-log phase (OD₆₀₀ ∼ 0.4) in the presence of 25 µg/mL chloramphenicol and 2 mM CaCl₂. In four separate tubes having 2 mL of LB and 2 mM CaCl₂, bacterial cultures were added to obtain the following percentages of defense-containing cells: 0%, 1%, 10%, or 100%. Cultures were inoculated with 20 µL of T3 or T5 phage stock and incubated overnight at 37°C with shaking. The tubes were centrifuged at 5,000 × g for 10 min, and 20 µL of the clarified supernatants were transferred into fresh cultures prepared under the same conditions. This passaging step was repeated for four consecutive rounds. Supernatants from the fourth passage were filter-sterilized (0.22 µm), serially diluted, and plated for plaque assays on defense-containing cells. Phage preparations that formed plaques were re-tested, and three individual plaques were picked for T3. No escapers could be isolated for T5. Each T3 plaque was used to inoculate 2 mL of mid-log defense-containing cultures (25 µg/mL chloramphenicol), which were grown overnight at 37°C with shaking and then centrifuged at 5,000 × g for 10 min at 4°C. The resulting supernatants were filter-sterilized (0.22 µm) and centrifuged at 113,000 × g for 2 h at 4°C. Pellets were gently resuspended in 300 µL of SM buffer, and final phage stocks were stored at 4°C.

Phage genomic DNA was extracted following previously described protocols^74^. Isolated phage stocks were incubated with DNase I and RNase A for 1 h to eliminate contaminating *E. coli* DNA. The nucleases were then inactivated by adding EDTA and heating at 65°C. Phage capsids were digested with proteinase K in the presence of SDS, and the resulting phage genomic DNA was purified by phenol–chloroform extraction. The purified DNA was sequenced using the Native Barcoding Kit V14 (Oxford Nanopore Technologies) on a GridION sequencer. Raw Nanopore reads were concatenated and assembled de novo using the Flye package^75^, and the assembled genome was compared to the WT T3 sequence.

### Bacterial growth assay

Chemically competent *E. coli* cells containing either WT or HNH-inactive retron (pACYCDuet-1 backbone) was transformed with WT or mutant exonuclease (pCDFDuet-1 backbone) and plated on LB agar having 2% glucose and grown at 37°C in the presence of appropriate antibiotics (chloramphenicol and streptomycin). Individual colonies were grown overnight in the presence of 2% glucose and antibiotics and 10-fold serial dilution of the liquid culture was spotted in LB agar plates containing either 2% glucose or 0.2 mM IPTG in the presence of antibiotics. The plates were incubated for 16 h at 37°C and imaged.

### msDNA cleavage analysis

Chemically competent *E. coli* cells containing the WT retron were transformed with WT or mutant T3 exonuclease and cultured overnight in LB media supplemented with 2% glucose and antibiotics (chloramphenicol and streptomycin). The overnight cultures were then used to inoculate a secondary culture and grown to OD_600_ of ∼0.4 in the presence of antibiotics. IPTG was added to induce expression of the exonuclease. DNA was extracted using the Zymo ZR Plasmid Miniprep kit and DNA shorter than 500 bps were resolved by 15% urea-PAGE.

### Fluorescence microscopy

Bacterial cells carrying either WT or HNH-inactive retron in the pACYCDuet-1 backbone were transformed with T3 exonuclease and grown in LB medium supplemented with the appropriate antibiotics (chloramphenicol and streptomycin) at 37°C until reaching an OD_600_ of 0.5–0.6. Cultures were induced with 0.2 mM IPTG for 2 h at 37°C with shaking. Following induction, cells were harvested by centrifugation and washed with 1× PBS. Cells were simultaneously incubated with 5 µg/mL Wheat Germ Agglutinin (WGA) Oregon Green (Invitrogen, W7024) and 10 µg/mL propidium iodide for 15 min at RT followed by two washes with 1× PBS. Cells were then incubated with 50 µg/mL 4′,6-diamidino-2-phenylindole (DAPI) for 10 minutes in the dark at room temperature. After staining, cells were transferred to Poly-L-lysine-treated coverslips and allowed to adhere for 5 minutes before imaging. Imaging was performed using an inverted spinning-disc confocal microscope built by 3i. The system included a Zeis 63x 1.4 NA Plan-Apochromat objective, Yokogawa CSU-W1 SORA confocal head, 2.8× magnification changer, 100 mW lasers (40% power: 405 nm, 488 nm, 561 nm), and a Hamamatsu Orca-Fusion BT camera (2x binning to create 74nm pixels). A piezo stage was used to acquire 10 z planes/sample with a step size of 0.27µm. Acquired images were analyzed using Fiji^76^. For presentation, only one z plane was used.

### tRNA sequencing and analysis

Chemically competent *E. coli* cells containing either the WT retron or the HNH-inactive (H71A) retron were transformed with T3 exonuclease and grown overnight in the presence of 2% glucose and antibiotics (chloramphenicol and streptomycin). Secondary cultures were inoculated from the overnight cultures, grown to an OD_600_ of 0.4–0.5 in the presence of antibiotics, and induced with IPTG for 15 min. Total RNA, including small RNAs, was extracted using the Zymo Quick-RNA Miniprep Kit.

tRNA sequencing (MSR-seq) was performed by MesoRNA (Chicago, IL, USA) as per previously reported protocol^77^. Total RNA (1 µg) was subjected to deacylation by oxidation with sodium periodate (NaIO₄), followed by quenching with ribose and treatment with sodium tetraborate for β-elimination. The deacylated RNA was then end-repaired using T4 polynucleotide kinase (PNK; New England Biolabs, NEB) and ligated to biotinylated capture hairpin oligos (CHO) containing sample-specific barcodes using T4 RNA Ligase I (NEB). After ligation, all barcoded samples were pooled. The pooled samples were captured on streptavidin-coated MyOne C1 Dynabeads (Thermo Fisher Scientific) via the biotinylated CHOs. Following capture, the samples were dephosphorylated with Quick CIP (Roche) and reverse-transcribed overnight using SuperScript IV Master Mix (Thermo Fisher Scientific). RNA templates were subsequently degraded with RNase H (NEB), and cDNA subjected to periodate oxidation. After quenching periodate with ribose, the single-stranded cDNAs were ligated to the second ligation oligo and then released from the beads by incubating at 95°C. For each sample, 100 ng of purified cDNA was PCR-amplified for 12 cycles (10 s at 98°C, 15 s at 55°C, 20 s at 72°C). The amplified products were purified using the DNA Clean & Concentrator kit (Zymo Research) and resolved on a 10% non-denaturing TBE polyacrylamide gel with appropriate dsDNA size markers. Bands corresponding to the final library size were gel-extracted and ethanol-precipitated. The resulting libraries were sequenced on a NovaSeq X Plus platform using a 1.5B flow cell (Illumina). Raw reads were demultiplexed and processed by MesoRNA to generate tRNA count tables. Reads were demultiplexed and trimmed in accordance with the MSR-seq library preparation protocol^77^. Quality control was performed with FastQC^78^. Reads were aligned to tRNA sequences in GtRNAdb^79^ using bowtie2^80^ to generate tRNA count tables.

tRNA sequencing counts were normalized and subjected to differential expression analysis using the DESeq2 package^81^ in R with custom scripts. Fold changes in tRNA abundance were calculated by DESeq2, which employs a negative binomial distribution model to estimate dispersion and determine statistical significance (p-value).

## Supporting information

Uncropped Images and Replicate Data

## Acknowledgements

We thank members of the B.W. laboratory for feedback and discussions, C. Hophan-Nichols and the cyber security team at Montana State University for computational support, C.M. Lawrence and Ravi Thakkar for maintenance and operation of the cryo-EM Core Facility at Montana State University. The cryo-EM Core Facility at Montana State University is supported by NSF no. 1828765 and the M. J. Murdock Charitable Trust. We thank Sam Mackintosh from IDeA National Resource for quantitative proteomics for mass spectrometry data. tRNA sequencing was performed by MesoRNA (Chicago, IL, USA). We thank Amy Binny Philip for the valuable suggestions on the statistics for tRNA sequencing analysis. Microscopy data was made possible with NIH grant (no. R35GM151262) to S.Z.D. Research in the Wiedenheft laboratory is supported by the National Institutes of Health (no. R35GM134867) and the Montana State University Agricultural Experimental Station (USDA NIFA). N.B. is supported by an F31 from National Institutes of Health (NIH) (no. GM153146) and received support from Montana INBRE (no. P20GM103474). Molecular graphics and analyses were performed with UCSF ChimeraX^82^, developed by the Resource for Biocomputing, Visualization and Informatics at the University of California, San Francisco, with support from NIH no. R01-GM129325 and the Office of Cyber Infrastructure and Computational Biology, National Institute of Allergy and Infectious Diseases. The funders had no role in the conceptualization, designing, data collection, analysis, decision to publish or preparation of the manuscript.

## Author Contributions

Project conceptualization was performed by B.W., R.A.W., J.T.G. and N.B. J.T.G, B.W. and N.B. wrote the manuscript with review and input from all authors. cyroEM data collection was performed by N.B. cryoEM data processing and structural analysis were generated by N.B. and J.T.G. Plaque assays were performed by S.D., H.L. and A.D. and A.G. Cloning was performed by J.T.G., R.A.W., S. D. and Q.M.P. Protein purification and biochemistry was performed by R.A.W. Bacterial toxicity assays were performed by J.T.G and Q.M.P. tRNA sequencing analysis was performed by J.T.G. Microscopy was performed by J.T.G. and S.Z.D. Covariation model for ncRNA and comparison between retron types was performed by M.B. Comparative analysis between *E. coli* and phage tRNAs were performed by J.T.G and M.B.

## Data Availability

Data for tRNA sequencing and phage whole genome sequencing are available in the National Center for Biotechnology Information (NCBI) Sequence Read Archive with BioProject ID: PRJNA1222438. Custom R script for tRNA sequencing analysis is available in GitHub ( https://github.com/WiedenheftLab/2025_George_Burman_et_al). EM maps of the type I-A retron complex and associated models were deposited to the Electron Microscopy Data Bank (EMDB) and Protein Databank (PDB) databases. Accession codes can be found in Supplementary Table 1. The PDB codes for the experimentally determined structure of the retron IA complex are 9N69, 9N6B, 9N6C. EMDB accession codes are EMD-49053, EMD-43103, EMD-49055 and EMD-49056. Raw micrograph images will be deposited to EMPIAR. All plasmids used in this study are available in Supplementary Table 2. Sequences for recombinantly expressed proteins and mass spectrometry data are available in Supplementary Table 3. Uncropped images and replicate data are available in Supplementary File 1. Datasets used and analyzed in this study can be obtained from the corresponding author on reasonable request.

---

**Supplementary Table 1:** Cryo-EM data collection, refinement and validation statistics.

|  | #1 Retron IA complex<br>“HNH Down”<br>(EMDB-49053)<br>(PDB 9N69) | #2 Retron IA complex<br>“HNH UP”<br>(EMDB-49055)<br>(PDB 9N6B) | #3 Retron IA complex<br>“without HNH”<br>(EMDB-49056)<br>(PDB 9N6C) |
| --- | --- | --- | --- |
| **Data collection and processing** |  |  |  |
| Magnification | 45,000 | 45,000 | 45,000 |
| Voltage (kV) | 200 | 200 | 200 |
| Electron exposure (e<sup>-</sup>/Å<sup>2</sup>) | 56.69 | 56.69 | 56.69 |
| Defocus range (μm) | -0.8, -2.5 | -0.8, -2.5 | -0.8, -2.5 |
| Pixel size (Å) | .9061 | .9061 | .9061 |
| Symmetry imposed | C1 | C1 | C1 |
| Initial particle images (no.) | 292,467 | 400,469 | 552,025 |
| Final particle images (no.) | 259,360 | 356,186 | 513,948 |
| Map resolution (Å) | 3.14 | 3.16 | 3.06 |
| FSC threshold | 0.143 | 0.143 | 0.143 |
| Map resolution range (Å) | 2.8-7.0 | 2.8-7.0 | 2.8-7.0 |
| **Refinement** |  |  |  |
| Initial model used (PDB code) | AlphaFold3 | AlphaFold3 | AlphaFold3 |
| Map sharpening methods | Phenix Half-Map | Phenix Half-Map | Phenix Half-map |
| Refinement Package | Phenix | Phenix | Phenix |
| Refinement Method | RealSpaceRefinement | RealSpaceRefinement | RealSpaceRefinement |
| **Model composition** |  |  |  |
| Non-hydrogen atoms | 20,417 | 20,489 | 17,299 |
| Protein residues | 2,445 | 2,442 | 2,107 |
| Nucleotide Residues | 114 | 114 | 114 |
| Ligands | 3 | 3 | 3 |
| **B factors (Å<sup>2</sup>)** |  |  |  |
| Protein | 48.6 | 37.54 | 33.69 |
| Nucleotide | 24.13 | 27.92 | 20.70 |
| Ligand | 83.85 | 71.64 | 56.02 |
| **R.m.s. deviations** |  |  |  |
| Bond lengths (Å) | 0.002 | 0.003 | 0.002 |
| Bond angles (°) | 0.459 | 0.544 | 0.518 |
| **Validation** |  |  |  |
| MolProbity score | 1.28 | 1.63 | 1.64 |
| Clashscore | 3 | 3 | 3 |
| Poor rotamers (%) | 1 | 0 | 2 |
| **Ramachandran plot** |  |  |  |
| Favored (%) | 96 | 96 | 95 |
| Allowed (%) | 4 | 4 | 5 |
| Disallowed (%) | 0 | 0 | 0 |

## Extended Data Figures

**Extended Data Figure 1.**
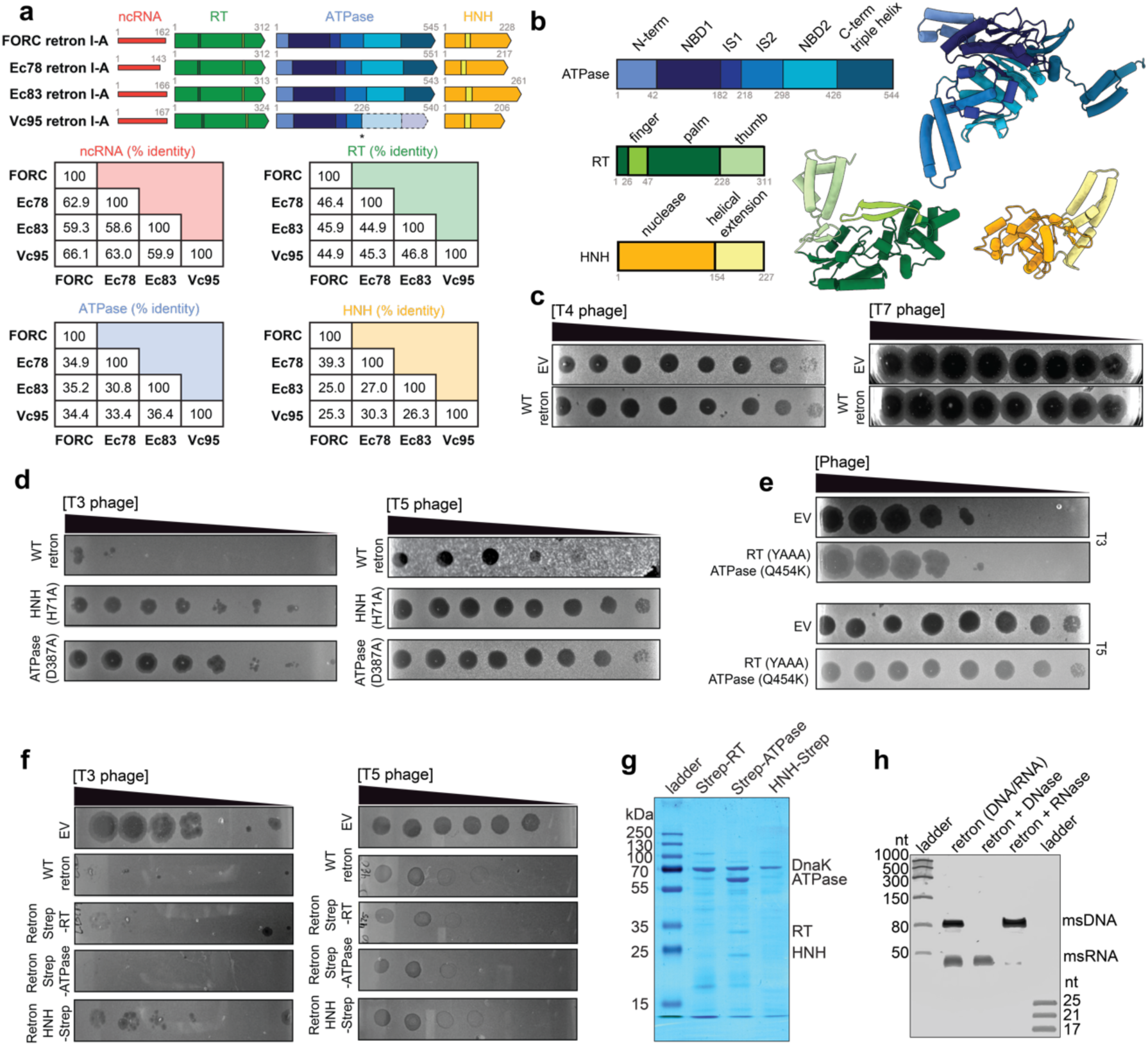
All components of the type I-A retron are necessary for phage defense. **a,** Sequence comparison of ncRNA, RT, ATPase and HNH between retron Ec78, Ec83, and Vc95 with the retron from *E. coli* FORC 82^14^. **b,** Domain architecture of the RT, ATPase and HNH nuclease mapped on AlphaFold3 predicted structures. **c**, Phage challenge assay with T4 and T7 phages in cells expressing either the type I-A retron immune system or an empty vector (EV) control. **d,** Mutations in the ATPase (D387A) or HNH nuclease (H71A) abolish retron-mediated immunity against T3 and T5. **e,** Compensatory ATPase mutation (Q454K) in retron-expressing RT mutant (YADD to YAAA) abolishes T3 and T5 immunity. **f,** An N-terminal Strep tag on the ATPase and RT does not affect retron-mediated immunity. In contrast, a C-terminal tag on the HNH nuclease partially impairs protection against T3, but does not affect T5 immunity. Data shown in panels c, d, e, and h are representative of *N = 2* biological replicates. **g,** Proteins purified using N-terminal tags on the RT and ATPase or C-terminal tags on the HNH were resolved using a 12% SDS-PAGE. **h,** Nucleic acids associated with the retron complex treated with either DNase or RNase and resolved using a 14% urea-PAGE.

**Extended Data Figure 2.**
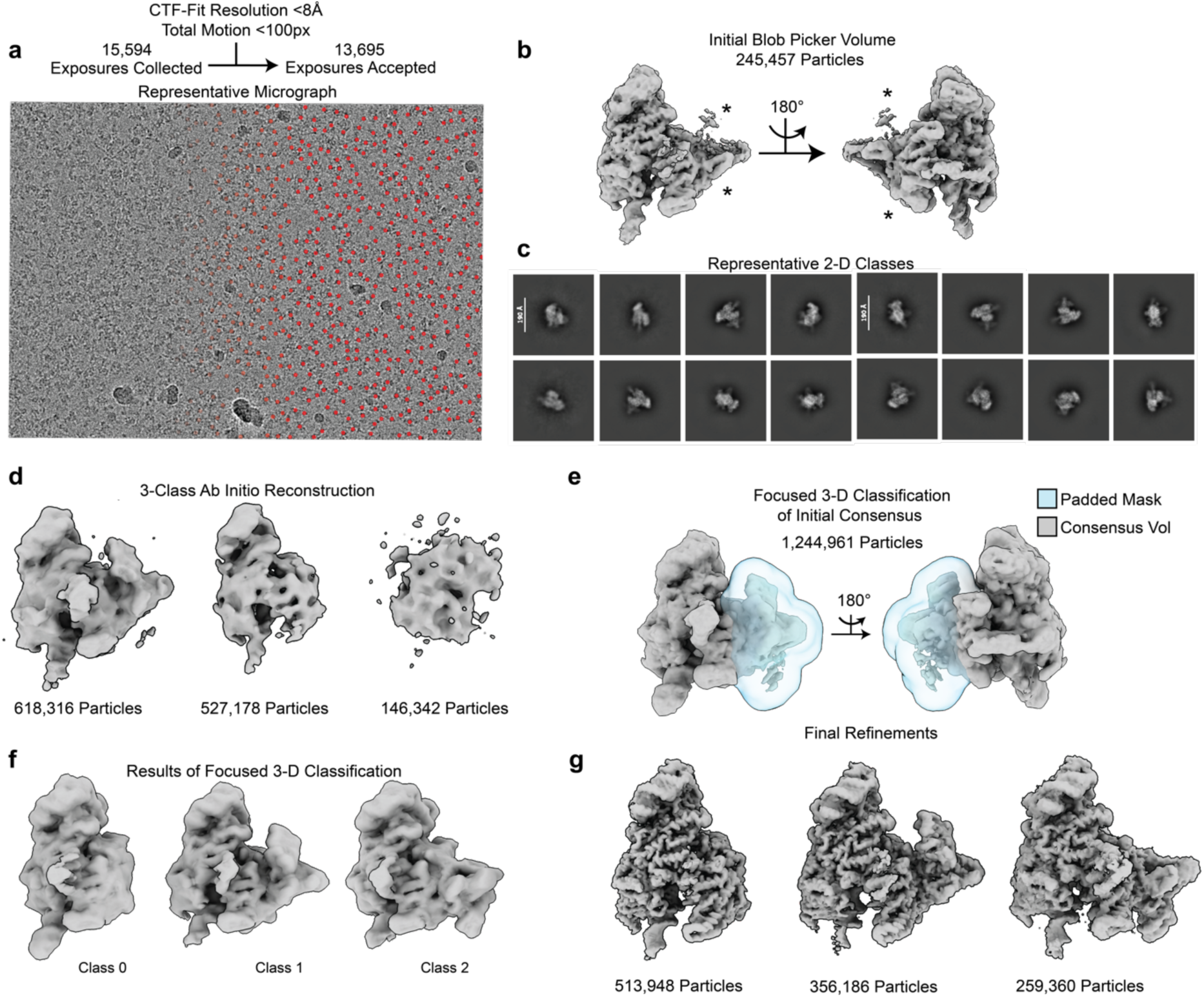
The workflow for cryoEM structure determination of the type I-A retron. **a**, Micrographs were filtered by contrast transfer function (CTF) and total frame motion before blob picking in cryoSPARC Live. **b**, *De novo* volume generated from the blob picker used for template picking. Template picks are shown as red dots right half of panel a. **c**, Representative 2D classes from template picked particles. **d**, 3-Class *ab initio* reconstruction used to remove junk from initial template picks. **e**, After re-extracting selected particles from panel d, focused 3D classification was used to parse particles based on HNH occupancy. **f**, Volumes obtained from focused 3D classification using the mask in panel e. Results are filtered to 8 Å. **g**, Final reconstructions of the type I-A retron complex.

**Extended Data Figure 3.**
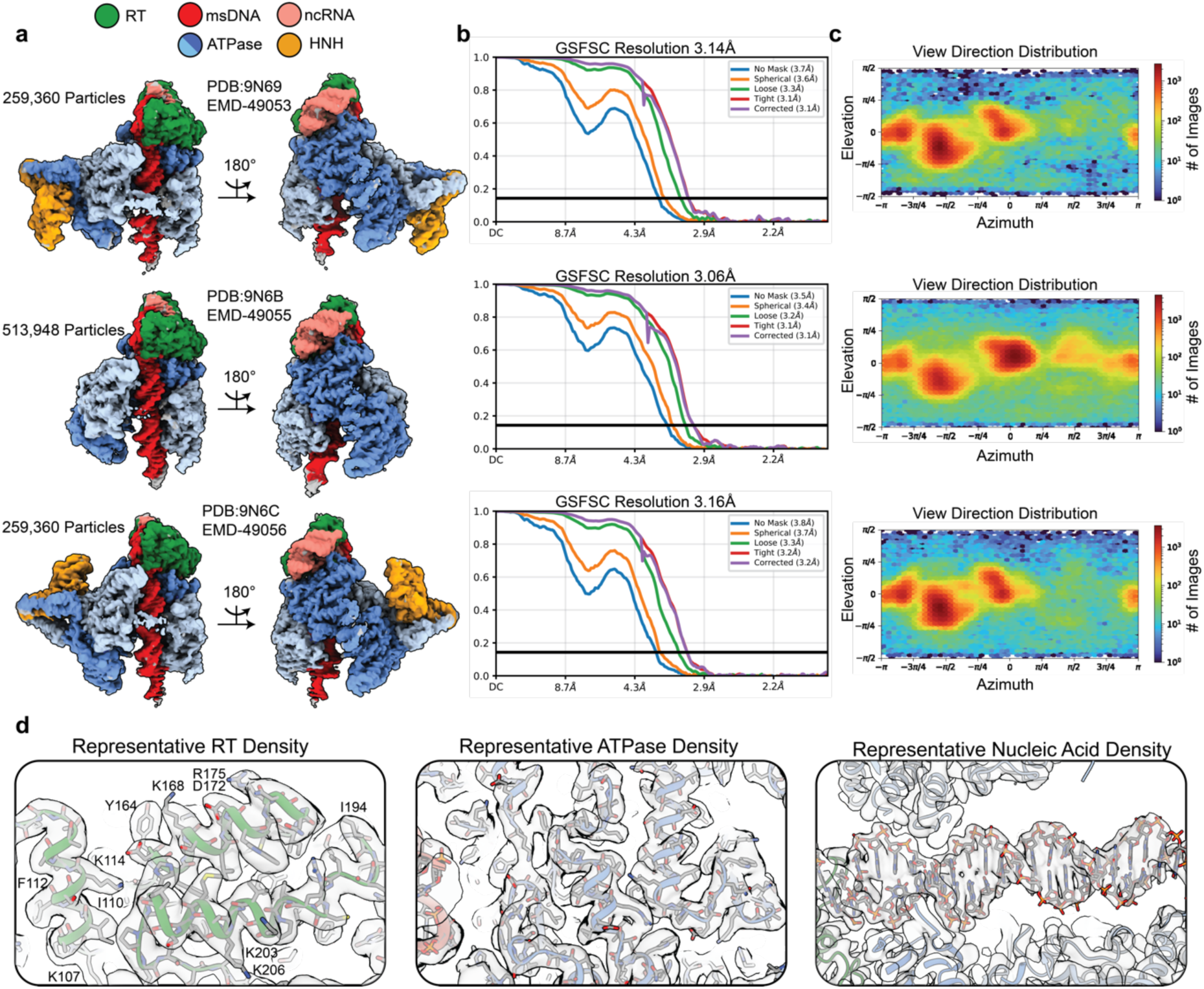
Structure of type I-A retron complex. **a,** Maps of the Retron IA complex. **b,** Gold Standard Fourier Shell Correlation from cryoSPARC were used to estimate resolution. **c,** Viewing distribution plots for each reconstruction reveals mild anisotropy. **d,** Close-ups demonstrating the model to map fit for different features of the complex including the RT, ATPase, and msDNA.

**Extended Data Figure 4.**
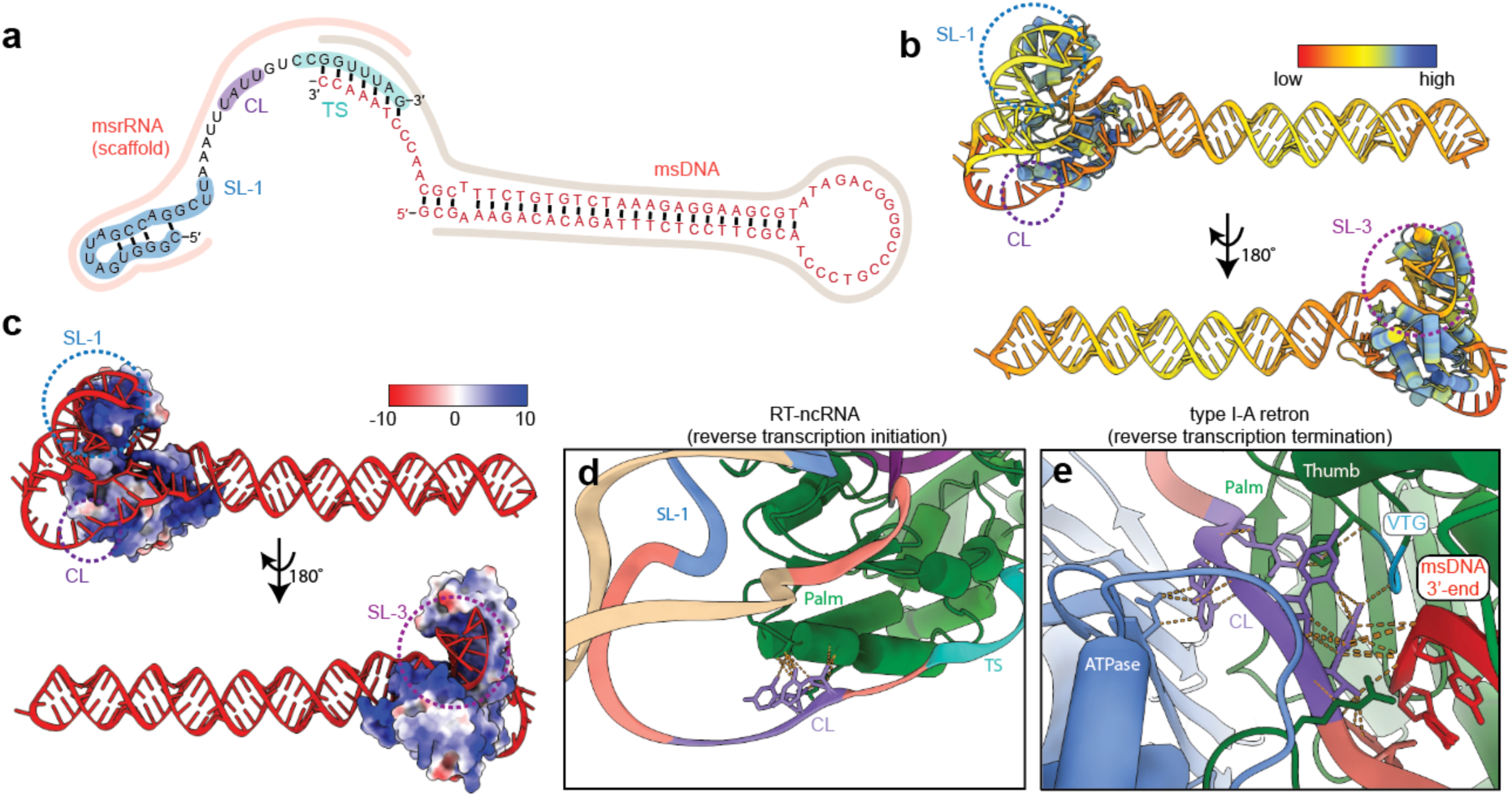
Comparison of the AlphaFold-predicted initiation complex and the experimentally determined post-assembly complex. **a,** The ncRNA sequence and msDNA sequence observed in the cryo-EM structure of retron I-A complex is mapped onto the covariance model shown in Fig. 1c. SL-1, CL and TS are indicated. **b,** AlphaFold3 prediction of the ncRNA-RT coloured by pLDDT confidence. **c,** The RT is coloured by electrostatics in the ncRNA-RT structure prediction. **d-e**, Cryo-EM structure of the surveillance complex reveals that the clamp (CL) interacts with the ATPase, 3′ end of the msDNA and RT (panel e). In contrast, the ncRNA-RT structure prediction shows CL engaged with the palm domain of the RT (panel d).

**Extended Data Figure 5.**
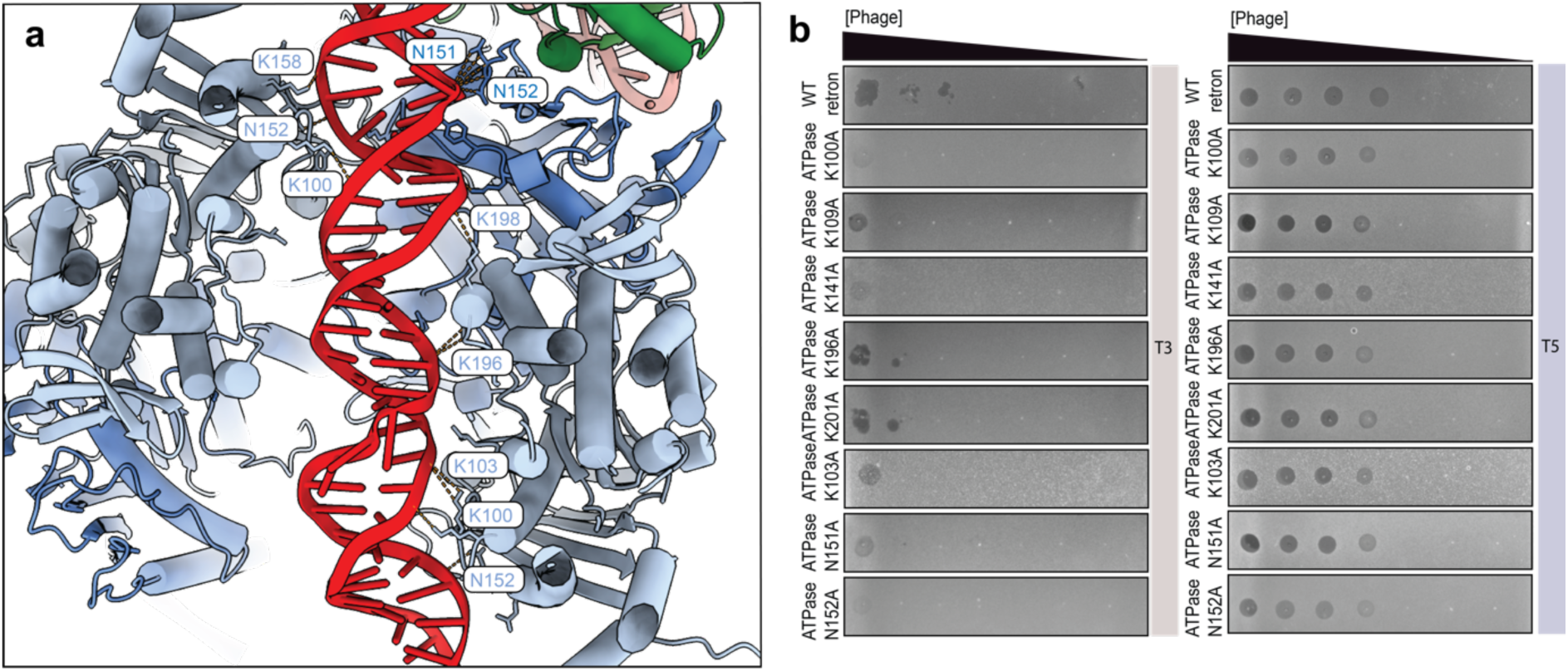
Positively charged residues of the ATPases are necessary for phage defense. **a,** Side chains of the ATPases that interact with the msDNA are highlighted. **b,** Point mutations in ATPase residues that interact with msDNA do not interfere with phage defense. Data shown in panels b is representative of *N = 2* replicates.

**Extended Data Figure 6.**
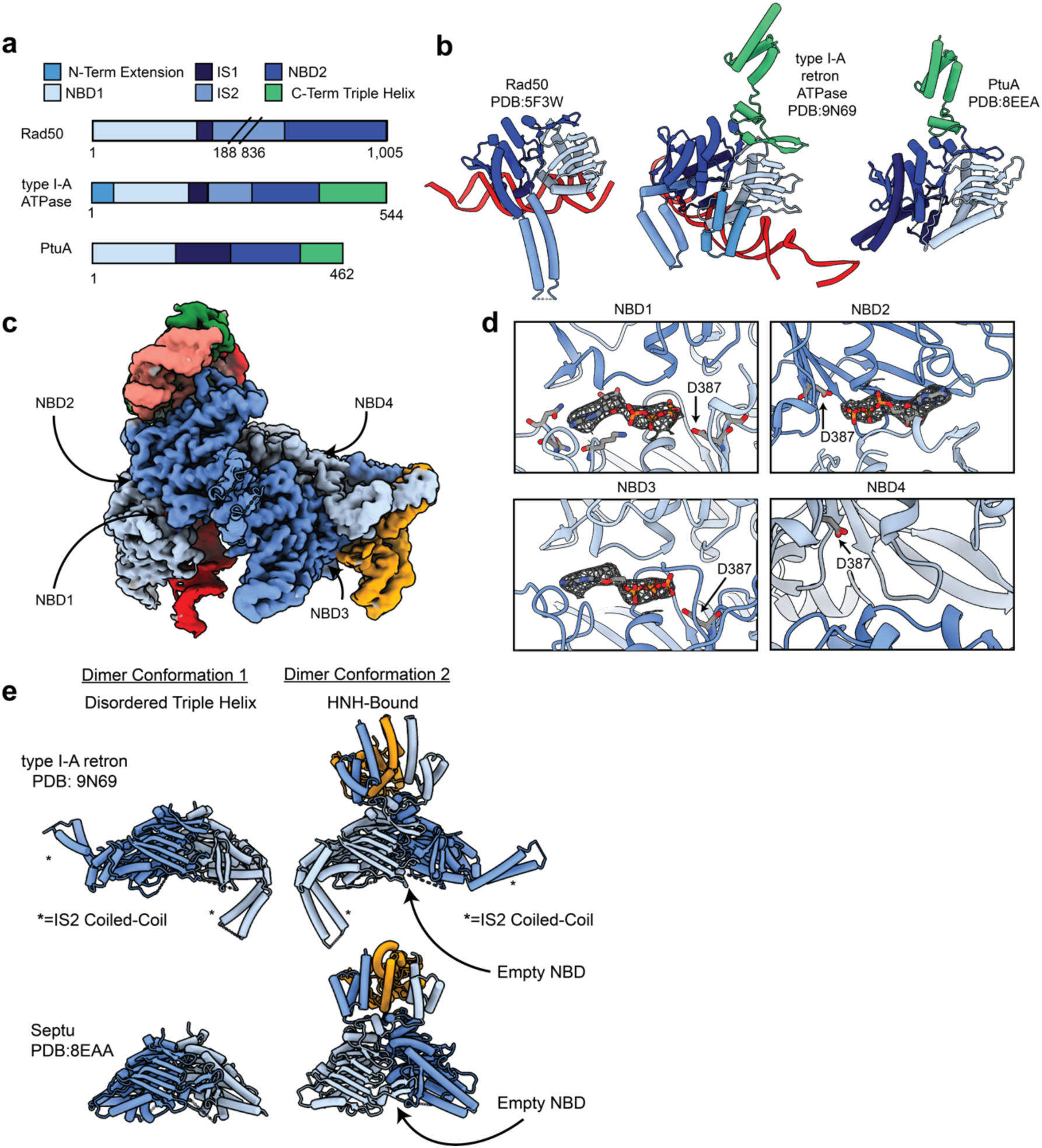
Structural features of the type I-A retron ATPase. **a,** Domain architecture of related SMC-family ATPases Rad50, Septu ATPase and type I-A retron ATPase^43,44^. **b,** Individual ATPase subunits from Rad50, retron, or Septu complexes are coloured according to domain architecture^43,44^. Retron ATPases contains a coiled-coil domain that wraps around DNA, similar to Rad 50^43^. However, in the retron, the coiled-coil domains from opposing homodimers provide a platform for interdimer interactions. **c,** The retron ATPases contains four NBDs **d,** In all experimentally determined retron structures, NBD4 is empty and has a binding pocket that is incompatible with ATP-binding **e,** Similar to Septu^44^, retron ATPase homodimers adopt two distinct conformations, one of which has a disordered C-terminal domain. However, unlike Septu, the retron ATPase forms a tetramer that is coordinated in part by the IS2 domains.

**Extended Data Figure 7.**
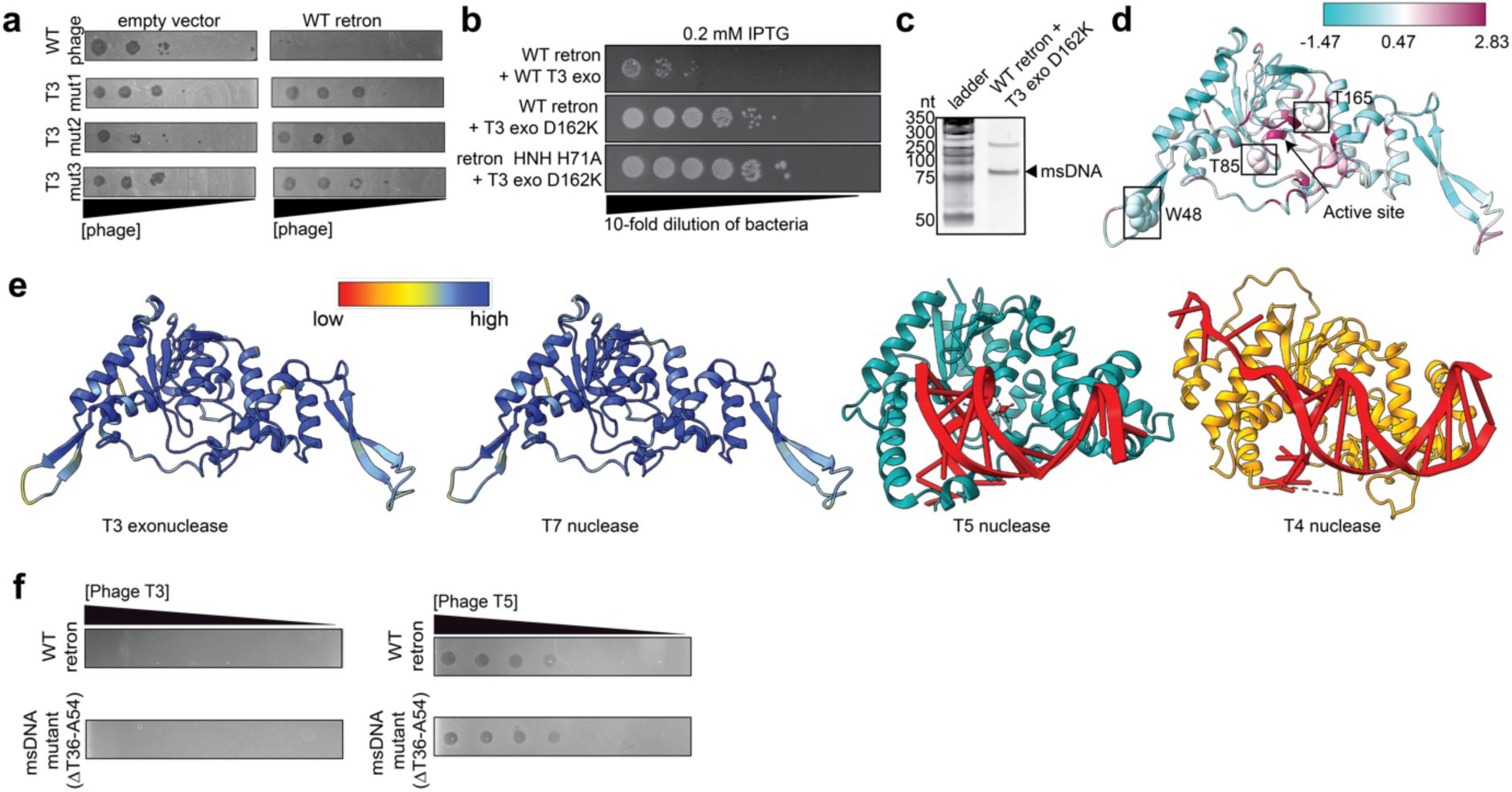
T3 exonuclease triggers retron immune system. **a,** Phage escaper mutants of T3 evade retron immunity. Empty vector (EV) control on the right, cells expressing the WT retron on the (right). **b,** Dilution series of *E. coli* cells expressing WT retron and mutant T3 exonuclease (D162K) shows lack of toxicity. **c,** Urea-PAGE reveals msDNA remains intact in cells expressing WT retron and mutant T3 exonuclease (D162K). Data shown is representative of *N = 2* independent biological replicates. **d,** Phage escape mutants are indicated on an T3 exonuclease structure prediction. Conserved residues are shown in violet. **e,** AlphaFold3 prediction for nuclease from T3 and T7, coloured according to pLDDT score. Cystal structures of related T5 (PDB: 5HNK) and T4 nucleases (PDB: 2IHN) are shown. **f,** Genetic truncation of the msDNA loop region from T35 to A54 does not abolish retron-mediated immunity. Data shown is representative of *N = 2* replicates.

**Extended Data Figure 8.**
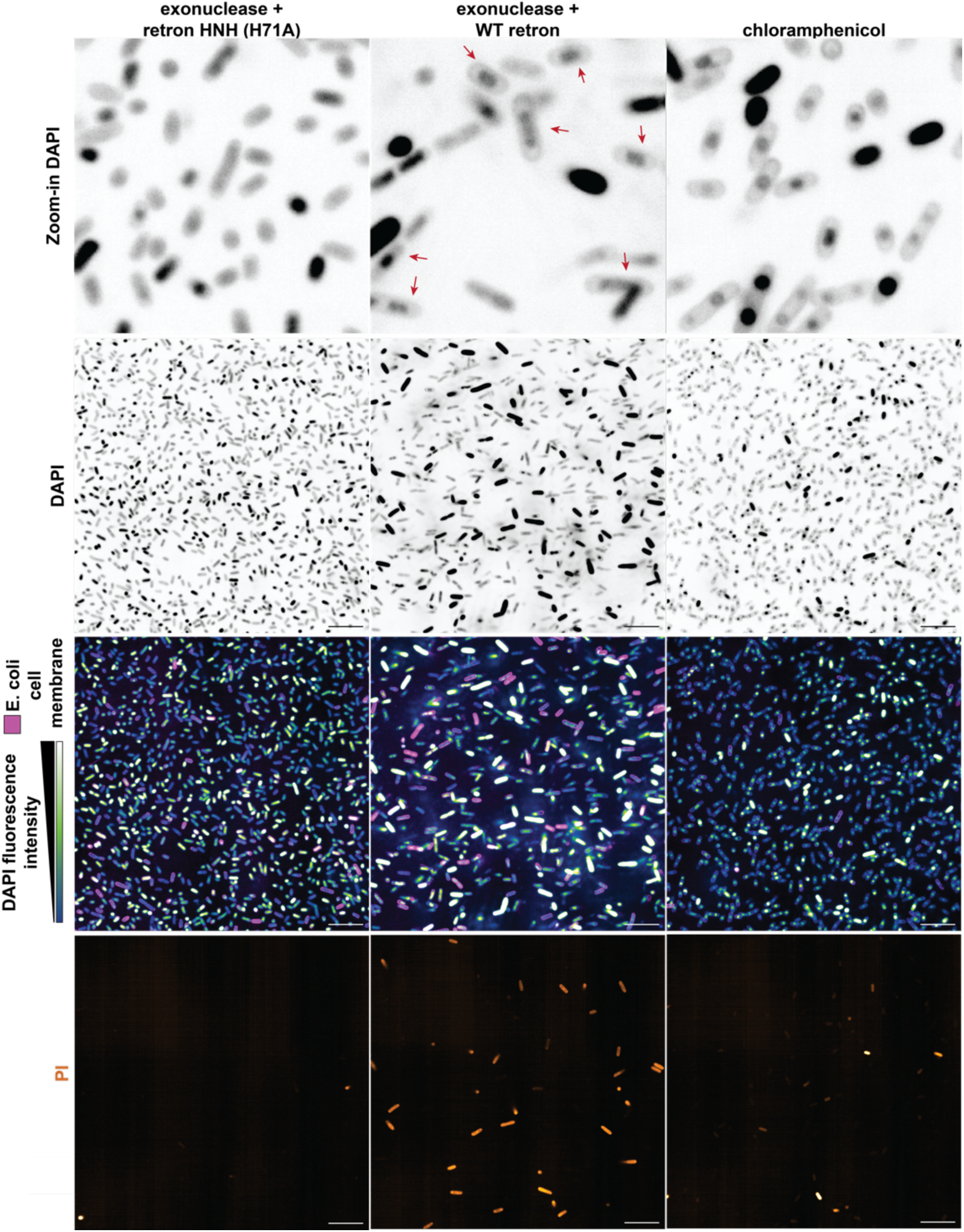
Retron activation leads to translational arrest. Fluorescence microscopy images of cells expressing either WT or HNH-inactive (H71A) retron with T3 exonuclease. Chloramphenicol treatment was used as a positive control for translational arrest. DNA is labeled with DAPI, the cell membrane with WGA-Oregon, and dead cells stained with propidium iodide. DAPI signal is shown in either pseudocolour or inverted grayscale. Red arrows indicate cells displaying nucleoid compaction. The scale bar represents 10 µm. Images representative of *N = 2* biological replicates.

**Extended Data Figure 9.**
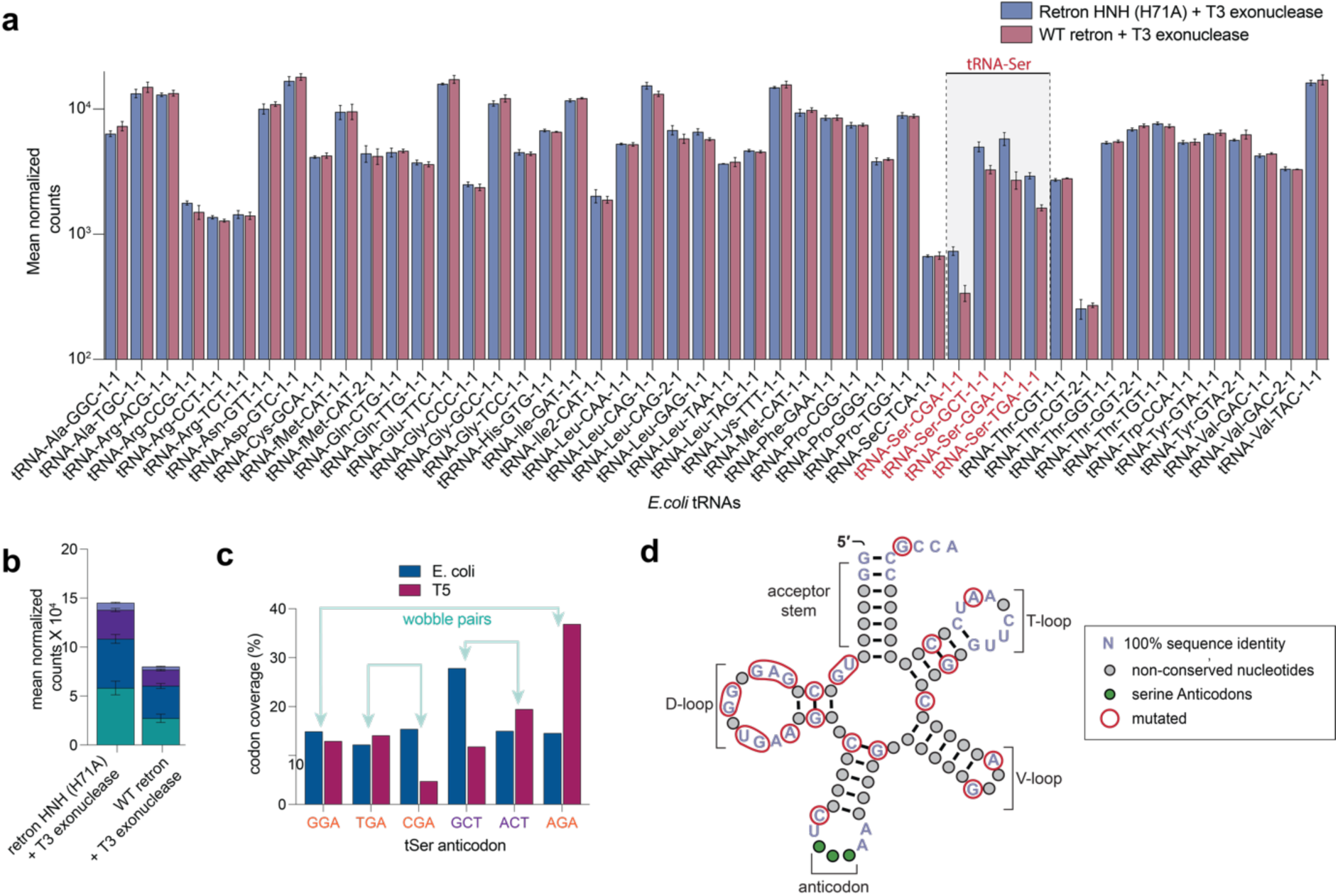
Type I-A retrons recognizes the D-loop of tRNAs for cleavage. **a,** Mean normalized counts of tRNA levels in *E. coli* cells expressing T3 exonuclease with either WT retron or HNH-inactive (H71A) retron. tRNA_Ser_ is highlighted in red. **b,** Normalized reads for total tRNA_Ser_ in cells expressing T3 exonuclease with either the WT retron or an HNH-inactive retron, as determined by tRNA sequencing. Data shown in panels a and b represents *N* = 3 independent biological replicates. **c,** Codon abundance in *E. coli* and WT T5 are shown. The codons decoded by phage tRNA_Ser_ in T5 are shown in orange, and codons not compensated by T5_MSU_ are highlighted in purple. **d,** Covariance model built from *E. coli* tRNA_Ser_ sequences. The regions conserved in *E. coli* tRNA_Ser_ and mutated in T5 tRNA_Ser_ are highlighted in red.

**Extended Data Figure 10.**
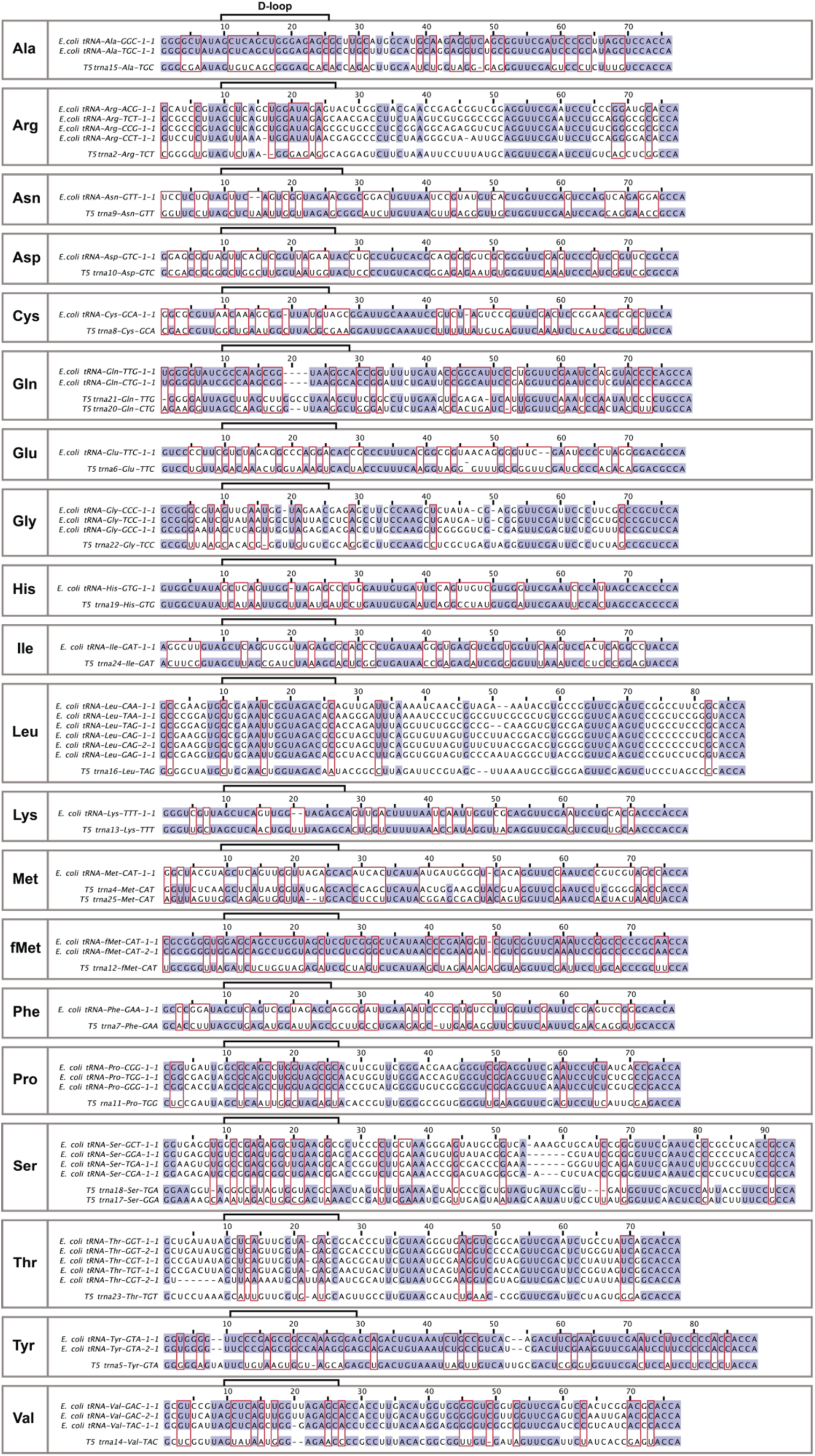
Alignment of *E. coli* tRNAs and WT T5 phage tRNAs. Conserved region are highlighted in lavender. Red boxes indicate residues conserved in *E. coli* tRNAs but different in in T5.

