## Supplementary figures and images for "Structural basis of antiphage defense by an ATPase-associated reverse transcriptase"

### Uncropped Images and Replicate Data

#### Uncropped Images and Replicate Data

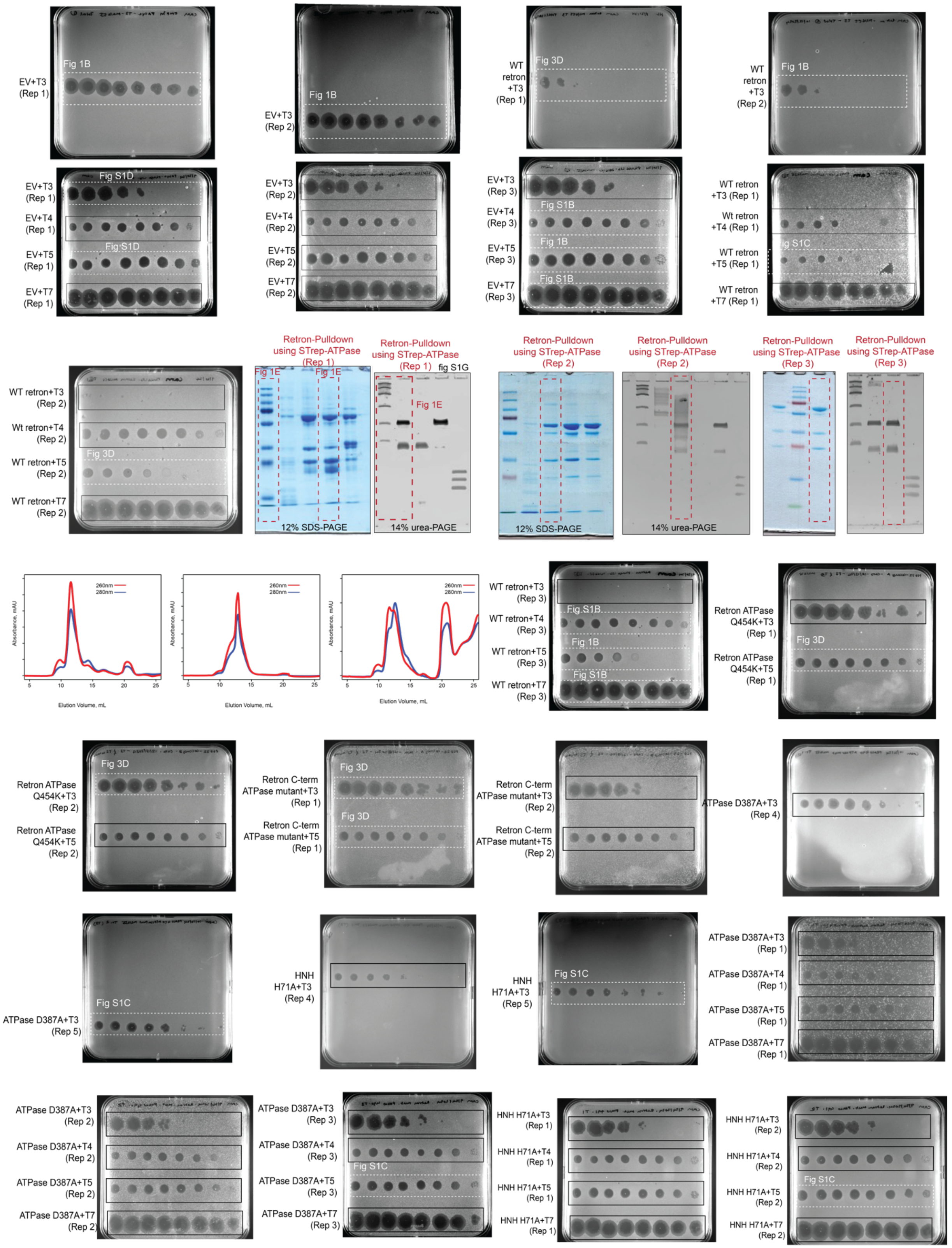

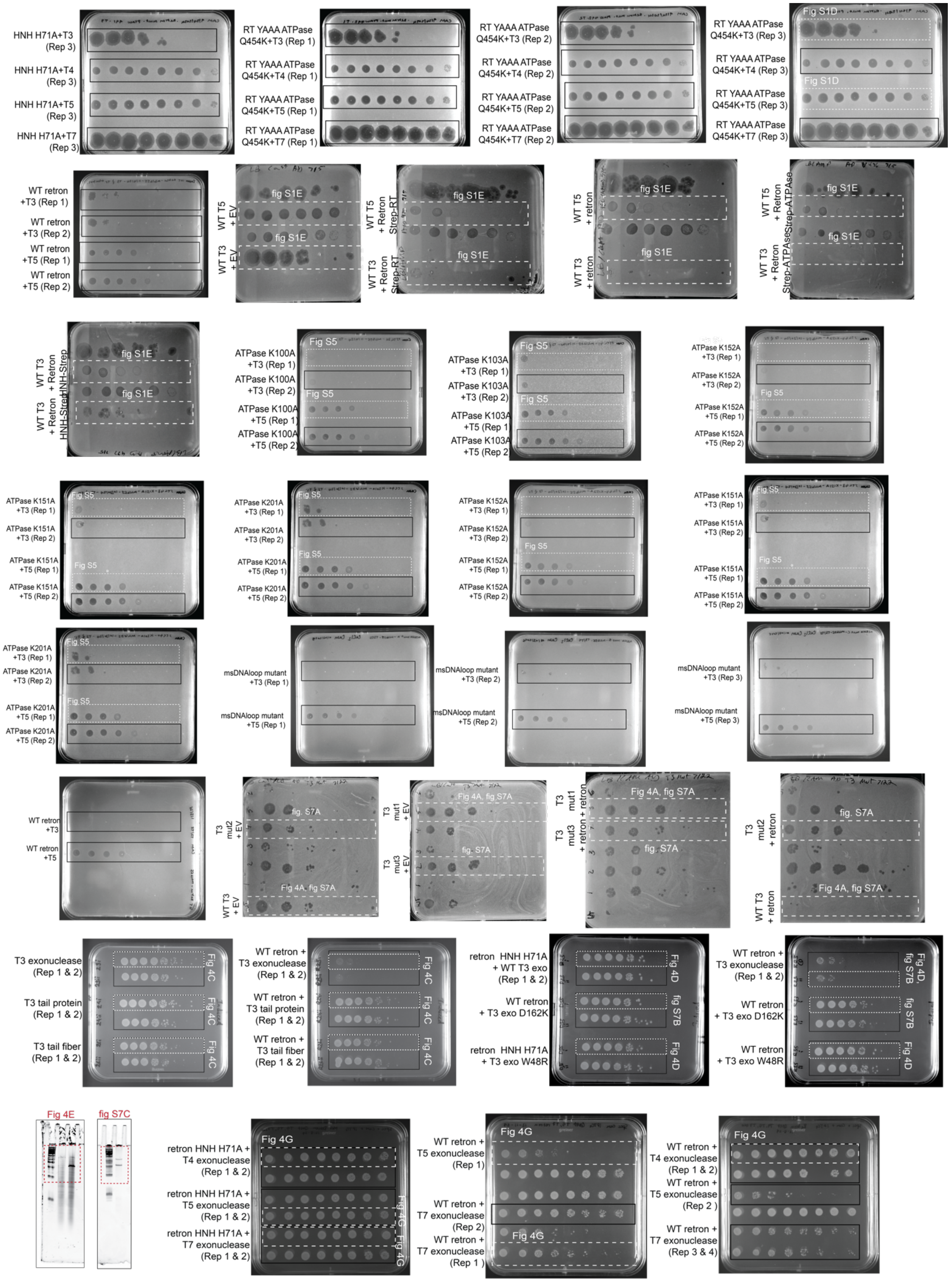

George, Burman *et al.* (2025)

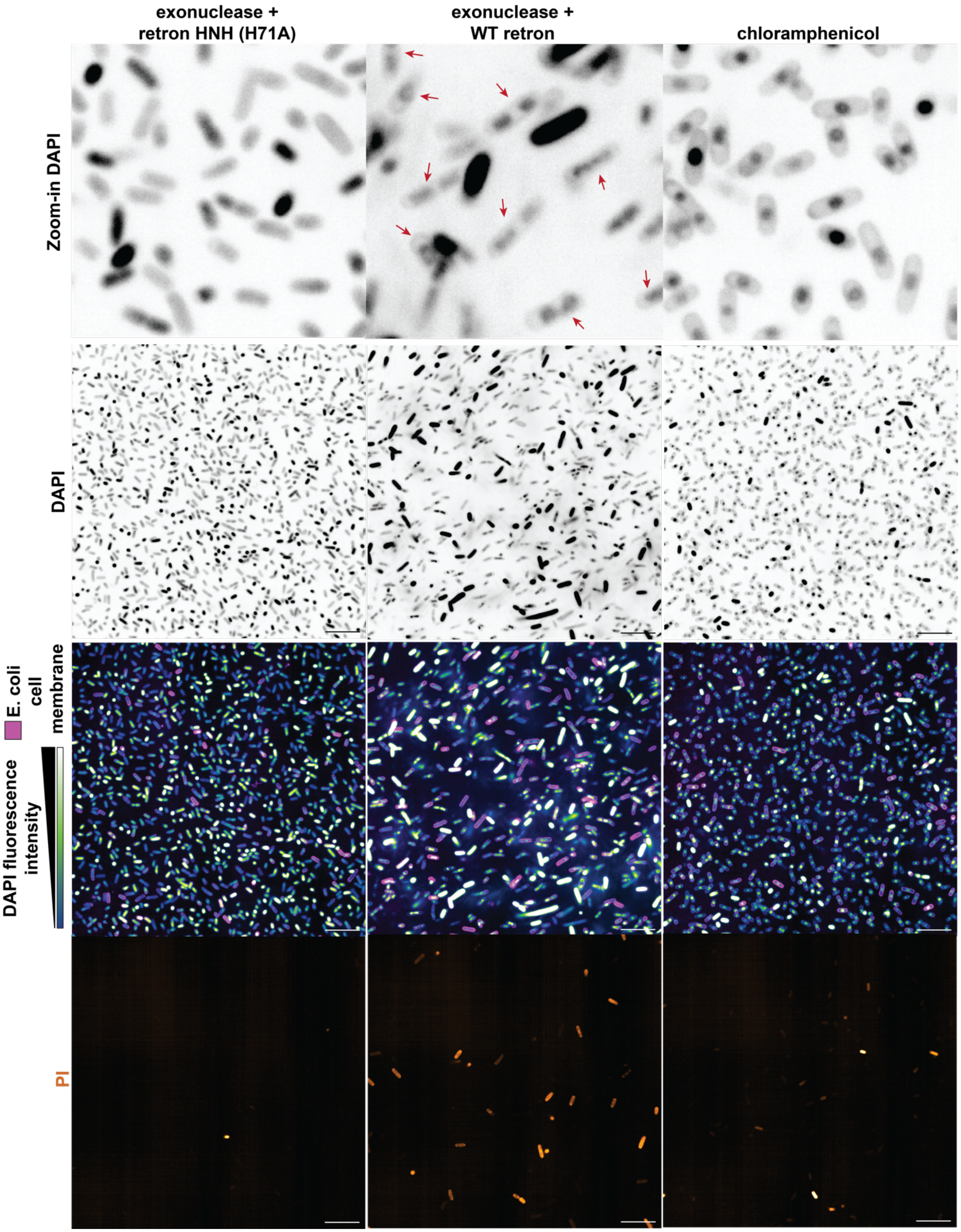
